# Harmonization of large multi-site imaging datasets: Application to 10,232 MRIs for the analysis of imaging patterns of structural brain change throughout the lifespan

**DOI:** 10.1101/784363

**Authors:** Raymond Pomponio, Guray Erus, Mohamad Habes, Jimit Doshi, Dhivya Srinivasan, Elizabeth Mamourian, Vishnu Bashyam, Yong Fan, Lenore J. Launer, Colin L. Masters, Paul Maruff, Chuanjun Zhuo, Ilya M. Nasrallah, Henry Völzke, Sterling C. Johnson, Jurgen Fripp, Nikolaos Koutsouleris, Theodore D. Satterthwaite, Daniel H. Wolf, Raquel Gur, Ruben Gur, John Morris, Marilyn S. Albert, Hans J. Grabe, Susan M. Resnick, R. Nick Bryan, David A. Wolk, Russell T. Shinohara, Haochang Shou, Christos Davatzikos

**Author notes:** for the ISTAGING Consortium, the Preclinical AD Consortium, the ADNI, and the CARDIA studies. Sharing senior authorship (Haochang Shou, Christos Davatzikos). Corresponding authors: Raymond Pomponio, Christos Davatzikos, 3700 Hamilton Walk, 7^th^ Floor, Center of Biomedical Image Computing and Analytics, University of Pennsylvania, Philadelphia, PA 19104, https://www.med.upenn.edu/cbica/.

## Abstract

As medical imaging enters its information era and presents rapidly increasing needs for big data analytics, robust pooling and harmonization of imaging data across diverse cohorts with varying acquisition protocols have become critical. We describe a comprehensive effort that merges and harmonizes a large-scale dataset of 10,232 structural brain MRI scans from participants without known neuropsychiatric disorder from 18 different studies that represent geographic diversity. We use this dataset and multi-atlas-based image processing methods to obtain a hierarchical partition of the brain from larger anatomical regions to individual cortical and deep structures and derive normative age trends of brain structure through the lifespan (3 to 96 years old). Critically, we present and validate a methodology for harmonizing this pooled dataset in the presence of nonlinear age trends. We provide a web-based visualization interface to generate and present the resulting age trends, enabling future studies of brain structure to compare their data with this normative reference of brain development and aging, and to examine deviations from normative ranges, potentially related to disease.

## 1. Introduction

Structural brain changes have been studied at various stages of the lifespan in relation to age and neurodegenerative diseases and conditions (Fjell and Walhovd, 2010; Habes et al., 2016), as well as to brain development (Courchesne et al., 2000; Sowell et al., 2001; Toga et al., 2006). A large number of imaging studies reported findings on age-related changes in brain structure during adolescence, early adulthood, and late adulthood (Giedd et al., 1999; Driscoll et al., 2009; Mills et al., 2016; Pfefferbaum et al., 1994; Tamnes et al., 2010; Terribilli et al., 2011). Traditionally, most neuroimaging studies have been limited to analyses on single-center homogeneous datasets carefully constructed using acquisition protocols that aim to minimize instrument-related variability in the data. However, in recent years there is an increasing trend towards data sharing in neuroimaging research communities, with multiple collaborative efforts for pooling existing data resources to form large, diverse samples covering a wide age range (Alfaro-Almagro et al., 2019; Thompson et al., 2014). Such collective efforts are critical for enabling development of diagnostic and prognostic biomarkers that apply across different imaging equipment as well as across the broad spectrum of demographics, which is essential for translation of neuroimaging research into clinical settings.

A number of studies have shown the importance of mega-analyses combining data from multiple cohorts. For example, data from the multi-site ENIGMA Consortium have been found to link volumetric abnormalities with post-traumatic stress disorder (Logue et al., 2018), schizophrenia (Van Erp et al., 2016), and major depressive disorder (Schmaal et al., 2016). However, there are important challenges in combining imaging data from multiple studies and sites. A major challenge is the lack of standardization in image acquisition protocols, scanner hardware, and software. Inter-scanner variability has been demonstrated to affect measurements obtained for downstream analysis such as voxel-based morphometry (Takao et al., 2011), lesion volumes (Shinohara et al., 2017), and DTI measurements (Zhu et al., 2011). Differences in sample demographics are also an important concern that should be handled carefully when combining multi-site data (LeWinn et al., 2017). For example, MR contrast may be confounded by differences in brain water content, which varies across age and diagnostic groups (Bansal et al., 2013). Finally, large-scale studies ultimately require robust and fully automated pipelines without the need to manually inspect and correct large sets of data, which is both time-consuming, subjective, and less likely to be adopted clinically.

In this paper we present a major effort designed to create the cross-sectional LIFESPAN dataset for quantitative characterization of structural changes in brain anatomy through the human lifespan from age 3 to 96. For this purpose, structural brain MRI scans from 18 studies were pooled together, creating a large, and most importantly, diverse sample (N=10,232). Although our focus is on structural MRI, our methodologies are applicable to any kind of imaging data. We test the robustness of a fully automated and standardized multi-atlas labeling pipeline, namely MUSE: *Multi-atlas region Segmentation utilizing Ensembles of registration algorithms and parameters and locally optimal atlas selection* (Doshi et al., 2016), which segments the brain into a set of hierarchically predefined regions of interest (ROIs) and measures the volume of each of these regions. A notable advantage of the multi-atlas segmentation methodology is that it computes the consensus labeling of a large ensemble of reference atlases, and hence simultaneously provides mechanisms for selecting atlases based on their local similarity to the target scan during the label fusion. The reference atlases represent anatomical variability across participants that span a wide age range, thus enabling a more robust segmentation across highly heterogeneous datasets.

The harmonization approach presented in this paper addresses the unique challenge of combining 18 studies from diverse age ranges in the presence of nonlinear age-related changes in brain volumes. Through the lifespan, the brain structure changes as a result of a complex interplay between multiple maturational and neurodegenerative processes. The effect of such processes could yield large spatial and temporal variations on the brain (Toga et al., 2006). A parsimonious model of age, such as a linear or quadratic model, is unlikely to sufficiently capture the relationship between age and volume throughout the lifespan (Fjell et al., 2010; Ziegler et al., 2012). Additionally, studies in our dataset did not overlap entirely on age, making techniques based on sample matching infeasible (Karayumak et al., 2019).

In order to capture non-linearities in age-related volume changes in brain anatomy through the lifespan, we propose to fit a generalized additive model (GAM) with a penalized nonlinear term to describe age effects (Hastie and Tibshirani, 1986; Wood, 2017). Within a single model, we estimated the location (mean) and scale (variance) differences in imaging measurements across sites. In the absence of ground truth, we performed simulation experiments to evaluate the harmonization performance across various conditions of sample composition. The simulation experiments leverage a large single-scanner study covering the entire adult lifespan to serve as an estimate of ground truth. Sampling this study and using simulations, we evaluate the effects of sample demographics and relative sample sizes on the harmonization accuracy.

Other communities that handle high dimensional data-integration across multiple sites have faced the necessity of harmonization. Among the available methods, ComBat, which was originally proposed to remove batch effects in genomics data (Johnson et al., 2007), has been recently adapted to diffusion tensor imaging data (Fortin et al., 2017), cortical thickness measurements (Fortin et al., 2018), and functional connectivity matrices (Yu et al., 2018). The method was shown to remove unwanted sources of variability, specifically site differences, while preserving variations due to other biologically-relevant covariates in the data. We adopt and test ComBat in our harmonization pipeline of the LIFESPAN dataset in conjunction with GAMs, which we refer to as ComBat-GAM. We compared ComBat-GAM to no harmonization and to ComBat with a linear model, based on their performances on a multi-variate brain age prediction task.

Successful harmonization of imaging measurements enabled us to estimate age-related volume changes for each anatomical region of the LIFESPAN dataset, which we refer to as age trajectories. The resulting age trajectories are supported by the large sample size of the dataset and may serve as a reference for the neuroimaging community. We provide an interactive online tool that will allow researchers to visualize the age trajectories of different anatomical regions, as well as to calibrate their own data with the LIFESPAN dataset, and position user-specific data among the reference trajectories.

## 2. Material and methods

### 2.1 MRI datasets

We collected structural MRI (T1) data from 18 studies. The pooled dataset included baseline scans of typically-developing and typically-aging participants from each study with available age and sex information. We defined typical development and typical aging as the absence of a known diagnosis of a neuropsychiatric disorder. We considered multi-center imaging studies that undertook efforts to unify protocols as single studies; this includes the Alzheimer’s Disease Neuroimaging Initiative (ADNI) (Jack Jr. et al., 2008), the Baltimore Longitudinal Study of Aging (BLSA) (Armstrong et al., 2019; Resnick et al., 2003), the Coronary Artery Risk Development in Young Adults study (CARDIA) (Friedman et al., 1988), the Pediatric Imaging, Neurocognition, and Genetics study (PING) (Jernigan et al., 2016), the Philadelphia Neurodevelopmental Cohort (PNC) (Satterthwaite et al., 2016), and UK Biobank (Alfaro-Almagro et al., 2019). Phases of ADNI (ADNI-1, ADNI-2) and BLSA (1.5T SPGR, 3T MPRAGE) were considered separate studies due to major scanner updates. A single scan was included in the LIFESPAN dataset for each ADNI and BLSA subject. Table 1 shows general characteristics of the study datasets. Figure 1 presents age distributions for each study in the LIFESPAN dataset, sorted by median age. Scanner models and acquisition protocol parameters in each dataset are given in Supplementary Table 1. Informed consent was obtained from all participants by the leading institutions of each individual study in the LIFESPAN dataset. The Ethics Committee of the leading institution of each cohort approved its study.

**Table 1.** Summary characteristics of the datasets included in the LIFESPAN dataset, sorted by median age.

| ***Dataset*** | ***Country of Origin*** | ***No. Participants*** | ***No. Females (%)*** | ***Age Range (Median)*** |
| --- | --- | --- | --- | --- |
| PING | USA | 301 | 151 (50.2) | [3, 21] (12.8) |
| PNC | USA | 1,422 | 742 (52.2) | [8, 24] (15.0) |
| MUNICH | Germany | 169 | 52 (30.8) | [18, 62] (29.7) |
| PennBBL | USA | 164 | 95 (57.9) | [16, 73] (30.2) |
| CHINA-TAH | China | 105 | 61 (58.1) | [20, 57] (32.0) |
| CARDIA | USA | 693 | 360 (51.9) | [42, 56] (51.0) |
| SHIP | Germany | 2,682 | 1,455 (54.3) | [21, 91] (52.8) |
| PAC-WASH | USA | 239 | 148 (61.9) | [42, 76] (61.5) |
| PAC-WISC | USA | 122 | 85 (69.7) | [48, 73] (62.2) |
| UKBIOBANK | United Kingdom | 2,151 | 1,159 (53.9) | [45, 80] (64) |
| PAC-JHU | USA | 92 | 56 (60.9) | [42, 88] (68.3) |
| BLSA-3T | USA | 935 | 501 (53.6) | [24, 96] (69.9) |
| ADC | USA | 103 | 66 (64.1) | [58, 95] (71) |
| AIBL | Australia | 429 | 242 (56.4) | [60, 92] (73) |
| ADNI-2 | USA | 312 | 170 (54.5) | [56, 95] (73.1) |
| BLSA-1.5T | USA | 91 | 35 (38.5) | [56, 86] (73.1) |
| PennPMC | USA | 40 | 21 (52.5) | [50, 85] (75) |
| ADNI-1 | USA | 182 | 84 (46.2) | [59, 89] (75.8) |
| **LIFESPAN dataset** |  | **10,232** | **5,483 (53.6)** | **[3, 96] (56.0)** |

**Figure 1:**
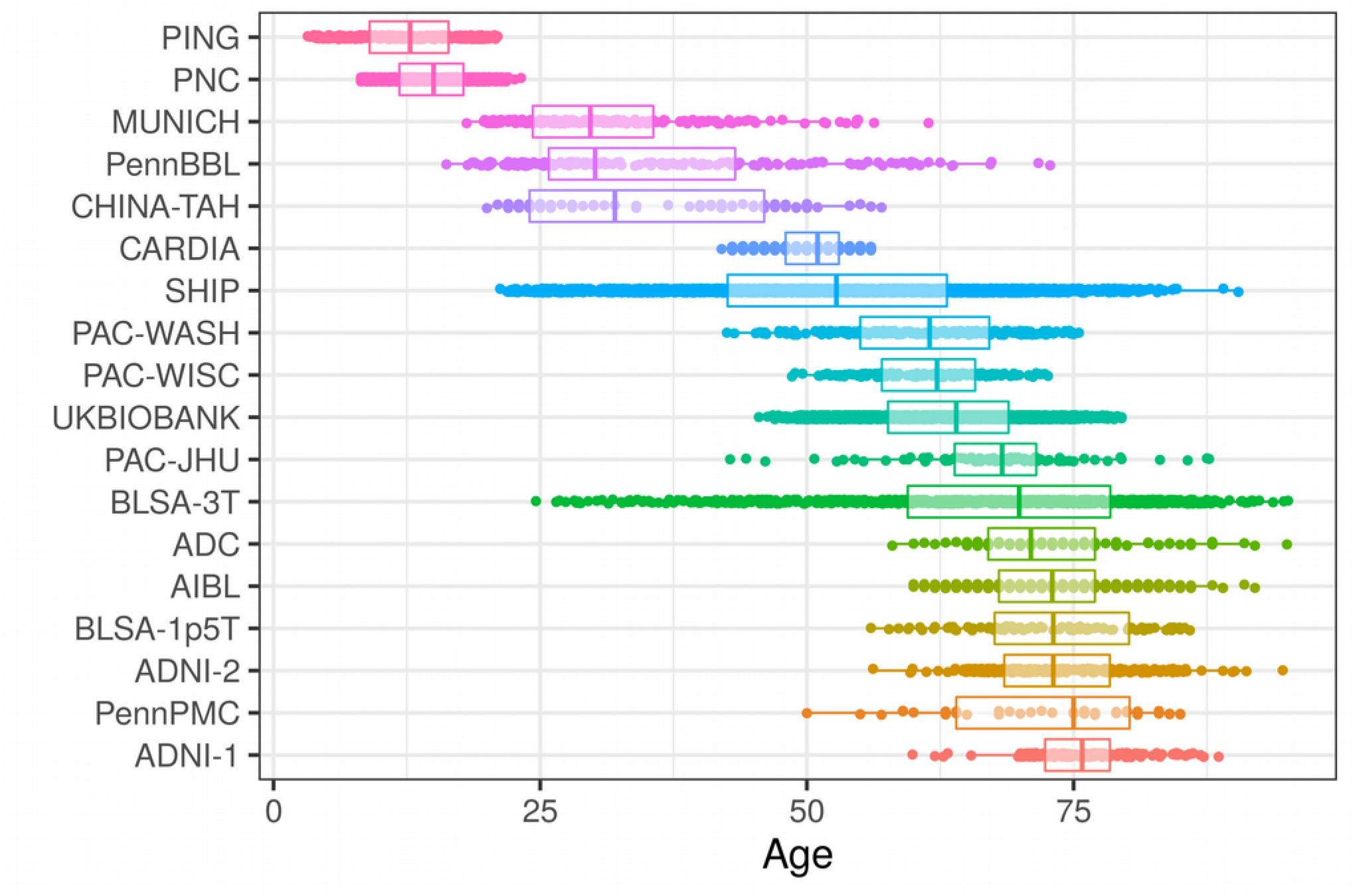
Age distributions of studies that are part of the LIFESPAN dataset, sorted by median age. The study with youngest median age, PING, contains participants from age 8 to 24. The study with oldest median age, ADNI-1, contains participants from age 59 to 89.

### 2.2 MRI image processing

A fully automated processing pipeline was applied to each participant’s T1-weighted scan. Pre-processing involved correction of magnetic field intensity inhomogeneity (Tustison et al., 2010) and skull-stripping, i.e. extraction of brain tissues, using a multi-atlas method (Doshi et al., 2013). For segmenting each T1 scan into a set of pre-defined anatomical regions of interest (ROIs) we used a multi-atlas, multi-warp label-fusion method, MUSE (Doshi et al., 2016), which has obtained top accuracy in comparison to multiple benchmark methods in independent evaluations (Asman et al., 2013). In this framework, multiple atlases with semi-automatically extracted ground-truth ROI labels are first warped individually to the target image using two different non-linear registration methods. A spatially adaptive weighted voting strategy is then applied to fuse the ensemble into a final segmentation. This procedure was used to segment each image into 145 ROIs spanning the entire brain. We calculated the volumes of these 145 ROIs, as well as the volumes of 113 composite ROIs that were obtained by combining individual ROIs into larger anatomical regions following a pre-defined ROI hierarchy. A list of the ROIs used in the LIFESPAN dataset is given in Supplementary Table 2.

### 2.3 Quality control of extracted variables

A systematic fully-automated quality control (QC) procedure guided by final outcome variables was conducted to identify and exclude cases of low quality. This procedure was applied on a set of 69 representative ROIs, including deep brain structures and sub-lobe level cortical parcellations, as well as the intra-cranial volume (ICV) (Supplementary Table 3). Volumes of selected ROIs were corrected for ICV and z-score transformed independently for each dataset in order to identify data outliers. We defined outliers as volumes that were greater than three standard deviations (SD) away from the within-study sample mean of the specific ROI. All scans that included at least three outlier ROIs were flagged for exclusion and investigation. In total, 254 scans were excluded based on this automated procedure, comprising 2.42% of the original sample (Supplementary Table 4).

### 2.4 Harmonization of imaging variables

We harmonized individual ROI volumes using a model that builds upon the statistical harmonization technique proposed in Johnson et al. (2007) for location and scale (L/S) adjustments to the data. This method estimates within a single model the location (mean) and scale (variance) differences in ROI volumes across sites, as well as variations due to other biologically-relevant covariates in the data that are intended to be preserved. Once estimated, the standardized ROI volumes can be achieved by removing location and scale effects due to site differences.

Given Y is the volume of an ROI, a LS adjustment of Y is:

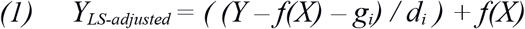

where *f(X)* denotes the variation of Y captured by the biologically-relevant covariates *X, g* is a vector of estimated location effects for each site *i*, and *d* is a vector of estimated scale effects for each site *i.* In the linear case, *f(X) = a + bX* and the corresponding adjustment is:

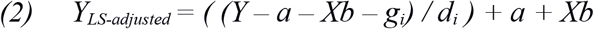

To allow for nonlinear age trajectories in ROI volumes informed by real data, we used a Generalized Additive Model (GAM) for *f(X)* in Equation 1. GAMs allow for flexible nonlinearity in *X* represented using a basis expansion. Additionally, penalization in the objective function of the model fitting ensures the smoothness of *f(X)* and avoids over-fitting to the observed data (Hastie and Tibshirani, 1986). In our design, we included a smoothed nonlinear term for age, using thin plate regression splines for basis expansion as described in Wood (2003), as well as parametric terms for sex and ICV. The model was estimated based on penalized regression splines and the degree of smoothness was internally selected using the restricted maximum likelihood (REML) criterion. Accordingly, our harmonized ROI volume was estimated as:

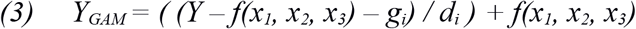

where *f*(*x*_*1*_, *x*_*2*_, *x*_*3*_) is the estimated function of the covariates age, sex, and ICV, respectively. In our model, we assumed that *f*(*x*_*1*_, *x*_*2*_, *x*_*3*_) was nonlinear in age, and linear in sex and ICV, hence:

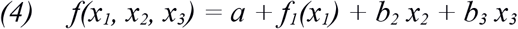

We integrated the non-linear GAM model with the previously-proposed framework of ComBat (Johnson et al., 2007) for the multivariate harmonization of multiple ROIs. The main premise of ComBat is that location and scale effects for multivariare outcomes, e.g. intensities across voxels or volumes across ROIs, are drawn from a common parametric prior distribution. ComBat estimates hyperparameters of the prior distributions from the data using empirical Bayes framework. Once estimated, the hyperparameters are used to compute conditional posterior estimates of all location and scale effects (Johnson et al., 2007). ComBat adjusts volumes Y using the conditional posterior estimates. Together with our non-linear GAM model, we have:

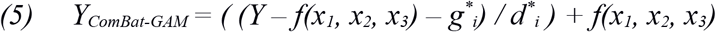

where *g*^*^ is a vector of conditional posterior estimates of location effects for each site *i*, and *d*^*^ is a vector of conditional posterior estimates of scale effects for each site *i*.

### 2.5 Evaluation of goodness of fit with GAM versus linear and quadratic models on single-site data

We first performed a comparative evaluation of the proposed GAM structure against both linear and quadratic models on single-site data. For the comparisons we selected three large studies with different age ranges. The Philadelphia Neurodevelopmental Cohort (PNC) included 1,422 participants from ages 8 to 24 (Satterthwaite et al., 2016). The Study of Health in Pomerania (SHIP) included 2,682 participants from ages 21 to 91 (Völzke et al., 2010). The 3-Tesla cohort of the Baltimore Longitudinal Study of Aging (BLSA-3T) included 935 participants from ages 24 to 96 (Armstrong et al., 2019). For each ROI, a linear model, a quadratic model, and a GAM with a smoothed nonlinear age term were fit to predict volumes from age. In all models, sex and ICV were included as additional covariates. The regression models were applied separately on each of the three study datasets to avoid confounding with site effects. We quantified the goodness of fit by calculating the adjusted R-squared for each model. Additionally, we used the Chi-square test to examine whether the residual sum of squares (RSS) were significantly lower using GAMs than other models.

### 2.6 Simulation experiments

The proposed harmonization model estimates a non-linear relationship between ROI volumes and age. The accuracy of the estimated age trajectory from multi-site data is a critical metric for harmonization performance. However, due to lack of ground-truth data, evaluations using real data were not possible. Therefore, we performed simulation experiments for assessing the effect of harmonization in the presence of known site effects for two different conditions. Toward this goal, we leveraged the large single-site SHIP study dataset (N=2,682) spanning ages 21 through 91.

In all experiments, we simulated volumes of the hippocampus for three hypothetical sites (named Site-A, Site-B and Site-C), using actual hippocampus volumes from SHIP. A ground truth age trajectory was first estimated on the entire SHIP data using a GAM model with a nonlinear term for age (sex and ICV effects were ignored for the simulations). For each of the 3 sites, we randomly sampled data following the sample size and age range constraints imposed by each experiment. Site-specific location and scale effects were then introduced on actual hippocampus volumes to generate the simulated datasets independently for each of the two experiments.

We performed harmonization using the LS adjustment with GAM method. The error of the estimated age trajectory after harmonization was quantified as the mean absolute error (MAE) over 100 equally-spaced age grid-points along the estimated trajectory versus the ground truth trajectory, standardized by the mean ROI volume, to produce relative Mean Absolute Error (rMAE). In all steps of the experiments, we repeated the simulation 100 times to obtain robust measures of estimated age trajectory error.

#### i. Effect of degree of overlap in the age ranges of data sites

Simulation Experiment I aimed to study the sensitivity of the proposed method to the amount of overlap in age ranges between harmonized datasets. For this purpose, we fixed the age ranges of Site-A and Site-C (30 to 50 and 60 to 80 years respectively), while allowing a 30-year sliding age range for Site-B that varies from younger (30 to 60 years) to older (50 to 80 years). All sites had equal sample sizes of 500.

#### ii. Effect of balancing sample sizes

Simulation Experiment II was designed to investigate harmonization of sites with unbalanced sample sizes. We assessed the effects of sub-sampling from a relatively larger site to create a balanced sample composition. Our main hypothesis was that leaving some data out of the harmonization in order to generate datasets balanced sample sizes might lead to more accurate alignment across studies. For this purpose, we fixed the sample size of sites A and C to 100, and varied the size of Site-B by randomly sub-sampling from 400 participants. We compared harmonization results using the complete Site-B sample (n=400) versus harmonization after sub-sampling Site-B by 50% (n=200) and 75% (n=100).

### 2.7 Harmonization of volumetric measurements from the LIFESPAN dataset

We applied ComBat-GAM on each of the 145 anatomical ROIs using the complete LIFESPAN sample to remove location and scale effects for each ROI.

Similar to Fortin et al. (2018), we evaluated the harmonization by assessing the accuracy on cross-validated brain age prediction using pre- and post-harmonized ROI volumes as features. The brain age model was constructed using a fully-connected neural network with one hidden layer, using the Keras and TensorFlow libraries (Abadi et al., 2015). ROI volumes for the complete LIFESPAN sample were used as input features to the model, in addition to sex and ICV. We performed 10-fold cross-validation, as well as leave-site-out cross validation to assess the effect of harmonization for brain age prediction on unseen sites. The network was trained with the Adam optimizer using mean squared error as the cost function with a constant learning rate of 1×10^−4^. The fully-connected layer consisted of 100 nodes with 20% dropout and RELU activation functions for each node. The output layer consisted of a single node with a linear activation function. We trained separate models with 10-fold cross validation on the complete LIFESPAN dataset using unharmonized ROIs, ROIs harmonized using ComBat with a linear model and ROIs harmonized using ComBat-GAM. The predictive accuracy of each model was evaluated using mean absolute error (MAE), i.e. mean absolute difference between predicted and actual ages. We also performed leave-site-out validations, using the PNC, SHIP, and BLSA-3T studies as independent test datasets in each experiment, in order to assess the effect of harmonization in predicting the brain age for data previously unseen by the training model.

### 2.8 LIFESPAN age trajectories of ROI volumes

After harmonization, we computed lifespan volumetric trajectories for each anatomical ROI, using GAM to model smoothed, nonlinear age trends. Since we were primarily interested in the relationship between age and ROI volumes, we regressed-out sex and ICV. The resulting age trajectories are free of sex and ICV effects, and enable a comprehensive analysis of brain volumes throughout the human lifespan.

Considering the large number of ROIs, we developed an interactive application that provides the end users a practical tool for selective visualization of the computed age trajectories for different brain regions. The visualization application, which allows users both to display LIFESPAN age trajectories and to position their own data after calibration with LIFESPAN data, was created with the Shiny package (Chang et al., 2019) in the R programming language, and is hosted at the following URL: https://rpomponio.shinyapps.io/neuro_lifespan/.

## 3. Results

### 2.5 Evaluation of goodness of fit with GAM versus linear and quadratic models on single-site data

Compared to linear models, GAMs achieved superior goodness-of-fit based on adjusted R-square for 124/145 ROIs in PNC, 120/145 ROIs in SHIP, and 128/145 ROIs in BLSA-3T. Compared to quadratic models, GAMs achieved superior goodness-of-fit based on adjusted R-square for 106/145 ROIs in the PNC, 114/145 ROIs in SHIP, and 129/145 ROIs in BLSA-3T. For most ROIs in all three studies, the Chi-square test indicated significant reduction in residual sum of squares (RSS) between linear models and GAMs (116/145 ROIs for SHIP, 94/145 ROIs for BLSA-3T, FDR corrected for multiple comparisons). A summary of the comparative evaluation is presented in Table 2.

**Table 2.** Results of evaluation of goodness of fit with GAM versus linear and quadratic models on single-site data.

|  | Number of ROIs in which GAM achieved better goodness of fit based on adjusted R-Square |  | Number of ROIs in which GAM achieved significant reduction in RSS based on Chi-square test* |  |
| --- | --- | --- | --- | --- |
| Study | GAM versus Linear | GAM versus Quadratic | GAM versus Linear | GAM versus Quadratic** |
| PNC (n=1422) | 124 (85.5%) | 106 (73.1%) | 88 (60.7%) | 0 (0%) |
| SHIP (n=2682) | 120 (82.7%) | 114 (78.6%) | 116 (80.0%) | 22 (16.3%) |
| BLSA-3T (n=935) | 128 (88.3%) | 129 (89.0%) | 94 (94.8%) | 12 (9.2%) |

*\*Chi-square test p-values were FDR corrected for multiple comparisons.*
*\*\*When comparing GAM to quadratic models with the Chi-square test, some p-values were not obtained due to estimated residual degrees of freedom from GAM being higher than residual degrees of freedom from a quadratic model. In such a case, GAM achieved a better fit than the quadratic model, while requiring fewer effective model parameters. Since it is not valid to perform the Chi-square test under such circumstances, we excluded these ROIs from this comparative experiment only. Specifically, 18 ROIs were excluded from PNC, 10 ROIs were excluded from SHIP, and 15 ROIs were excluded from BLSA-3T.*

Figure 2 presents hippocampus volumes in the three selected studies with separate fits using linear models, quadratic models, and GAMs.

**Figure 2:**
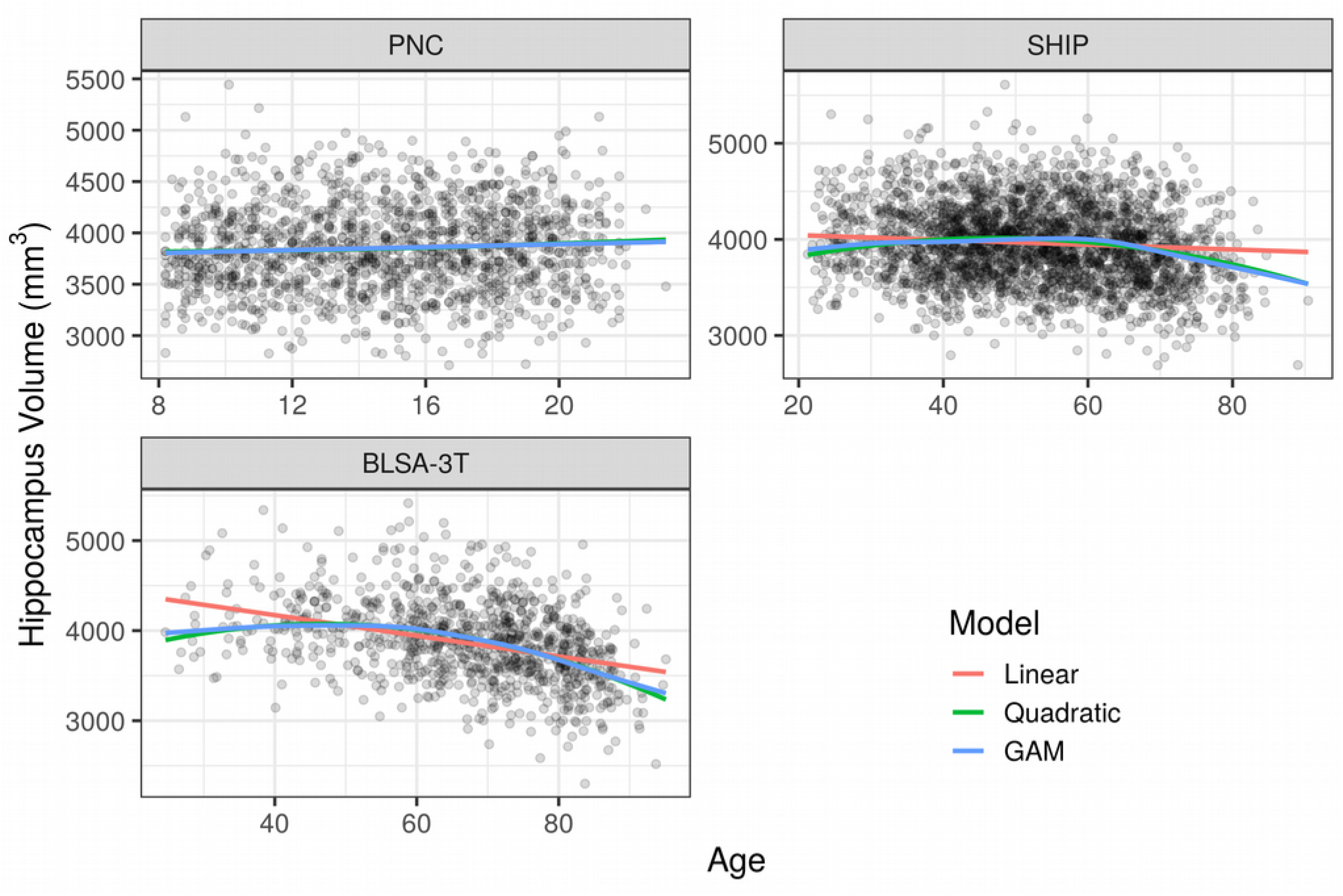
Comparison of age trajectory fits for the hippocampus volume from three studies (PNC, SHIP, and BLSA-3T) using linear models, quadratic models, and GAMs. The trajectories plotted are for females and assume an average intra-cranial volume (ICV). In the top-left panel, the difference between fits is not distinguishable. In the top-right panel and the bottom-left panel, both the quadratic fit and the GAM fit exhibit clear improvement over the linear fit.

### 3.2 Simulation experiments

#### i. Effect of degree of overlap in the age ranges of data sites

In Figure 3, we present four of the possible scenarios under the constraints of Simulation Experiment I, which assessed the impact of various degrees of overlap among three sites. The age range of Site-B had a fixed width but was free to vary from younger to older ages. The relative Mean Absolute Error (rMAE) of age trajectory estimation for different age ranges of Site-B, before and after harmonization, are shown in Figure 4. Age trajectories were more accurately estimated with harmonized data than without harmonization, with median rMAE decreasing from 0.033 to 0.015 when data were harmonized. With the harmonized data, the estimation error decreased consistently as the age range of Site-B moved from younger to older until reaching a plateau for the age range 45 to 75, after which estimation error began increasing with older age ranges.

**Figure 3:**
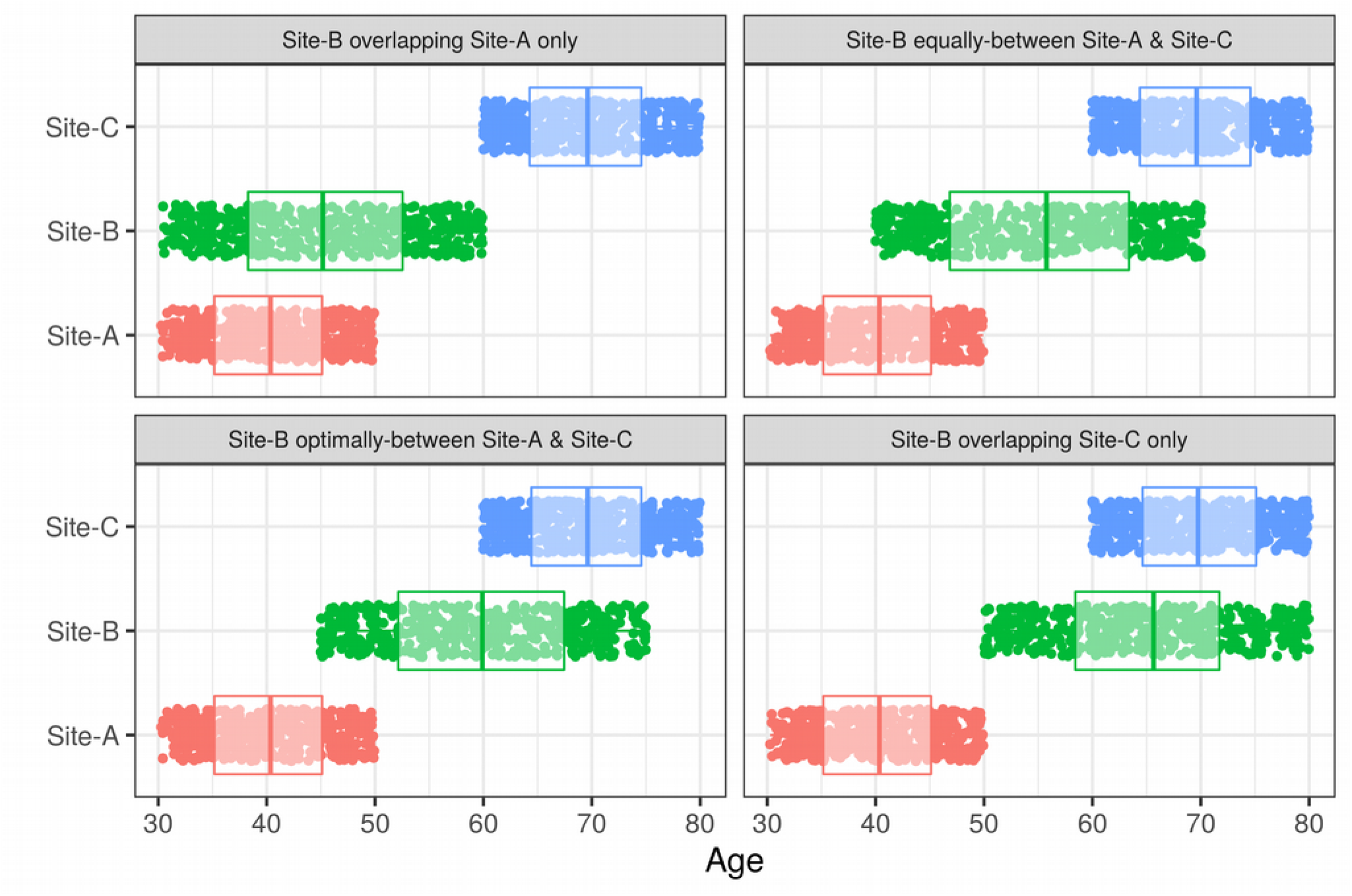
Four possible scenarios under the constraints of Simulation Experiment I, which assessed the effect of different degrees of age overlap on harmonization performance. The age range of Site-B was free to vary from younger to older ages. In the upper-left panel, Site-B is overlapping only Site-A and not Site-C. In the upper-right panel, Site-B is equally-overlapping Site-A and Site-C. In the lower-left panel, Site-B is optimally overlapping Site-A and Site-C based on the results of the experiment. In the lower-right panel, Site-B is overlapping only Site-C and not Site-A.

**Figure 4:**
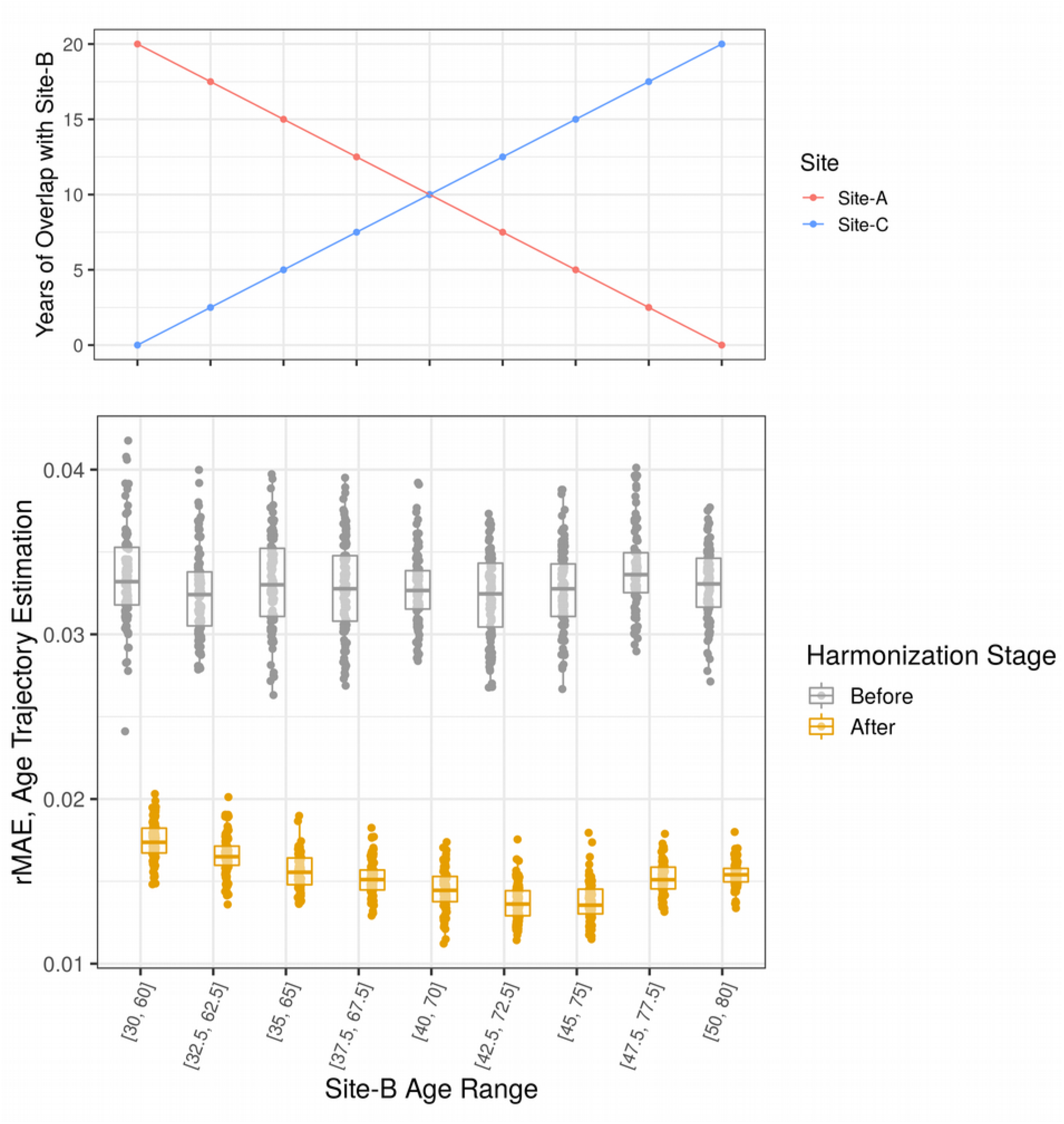
The relationship between the relative Mean Absolute Error (rMAE) of age trajectory estimation and the age range of Site-B in Simulation Experiment I. In both panels, the x-axis indicates the age range of Site-B, and moving from left to right is equivalent to sliding the age range of Site-B from younger to older ages. In the top panel, the proportion of age overlap between Site-B and each of the other two sites is plotted. In the bottom panel, the relative error in estimating the age trajectory is shown both before and after harmonization. Harmonization reduced the estimation error. Additionally, having partially-overlapping age ranges between the three sites led to reduced error, with a bias towards a larger overlap with the older Site-C rather than an equal overlap with Sites A and C.

These results suggest that a partial overlap of 5 to 10 years between consecutive dataset pairs is sufficient for optimal harmonization of multiple datasets, even if the combination of age ranges includes disjoint pairs. The LIFESPAN dataset included three studies (PNC, SHIP, UKBIOBANK) with sample sizes greater than 1,000 and age ranges spanning more than 15 years (Figure 1). These datasets played a key role in connecting other smaller datasets together via partially-overlapping age ranges. The weakest overlap was in the young-adult age range of 18 to 25 years. We included two studies (MUNICH, PennBBL) that covered this age range entirely and had sample sizes greater than 150, as well as three additional studies that partially-covered this age range (PING, PNC, CHINA-TAH).

#### ii. Effect of balancing sample sizes

Age trajectory estimation errors for varying amounts of sub-sampling from the relatively large site are shown in Figure 5. The optimal performance was achieved when all data points were used, even though the relative ratio of sample sizes between sites was heavily unbalanced (n=400 vs n=100). These results suggest that the negative impact of reduced sample sizes is greater than that of unbalanced sample compositions in age trajectory estimation after harmonization, which is in contrast to our original hypothesis that balanced datasets would lead to better harmonization.

**Figure 5:**
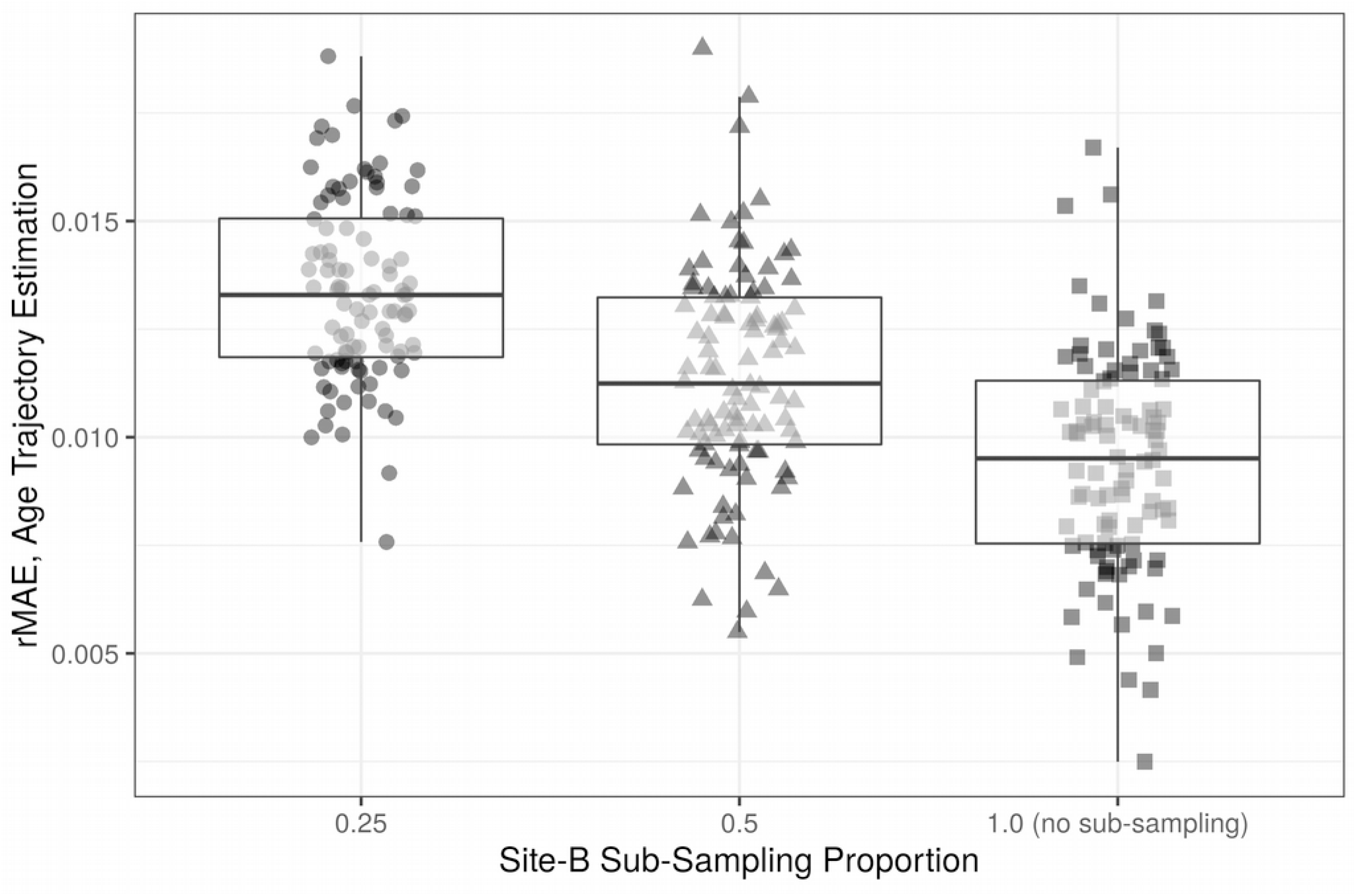
The relationship between the relative Mean Absolute Error (rMAE) of age trajectory estimation and the proportion of sub-sampling from Site-B in Simulation Experiment II. The original sample size of Site-B was four times larger than that of Sites A and C. At 0.25, the size of Site-B after sub-sampling was equal to the size of Sites A and C. At 0.5, the size of Site-B after sub-sampling was equal to the twice the size of Sites A and C. Results were optimal when all data points were used.

### 3.3 Harmonization of volumetric measurements from the LIFESPAN dataset

Our proposed harmonization method removed location and scale effects associated with site, after controlling for age, sex, and ICV with GAM. Figure 6 shows the adjustments made to hippocampus volumes after harmonization. Adjustments for other important structures, as well as for total gray matter and total white matter, are shown in Supplementary Figure 1. After harmonization, the residual volumes by site are centered at zero as expected, indicating the removal of location effects. Scale effects were not as strong for the hippocampus, with the residual volumes by site showing similar variances before and after harmonization.

**Figure 6:**
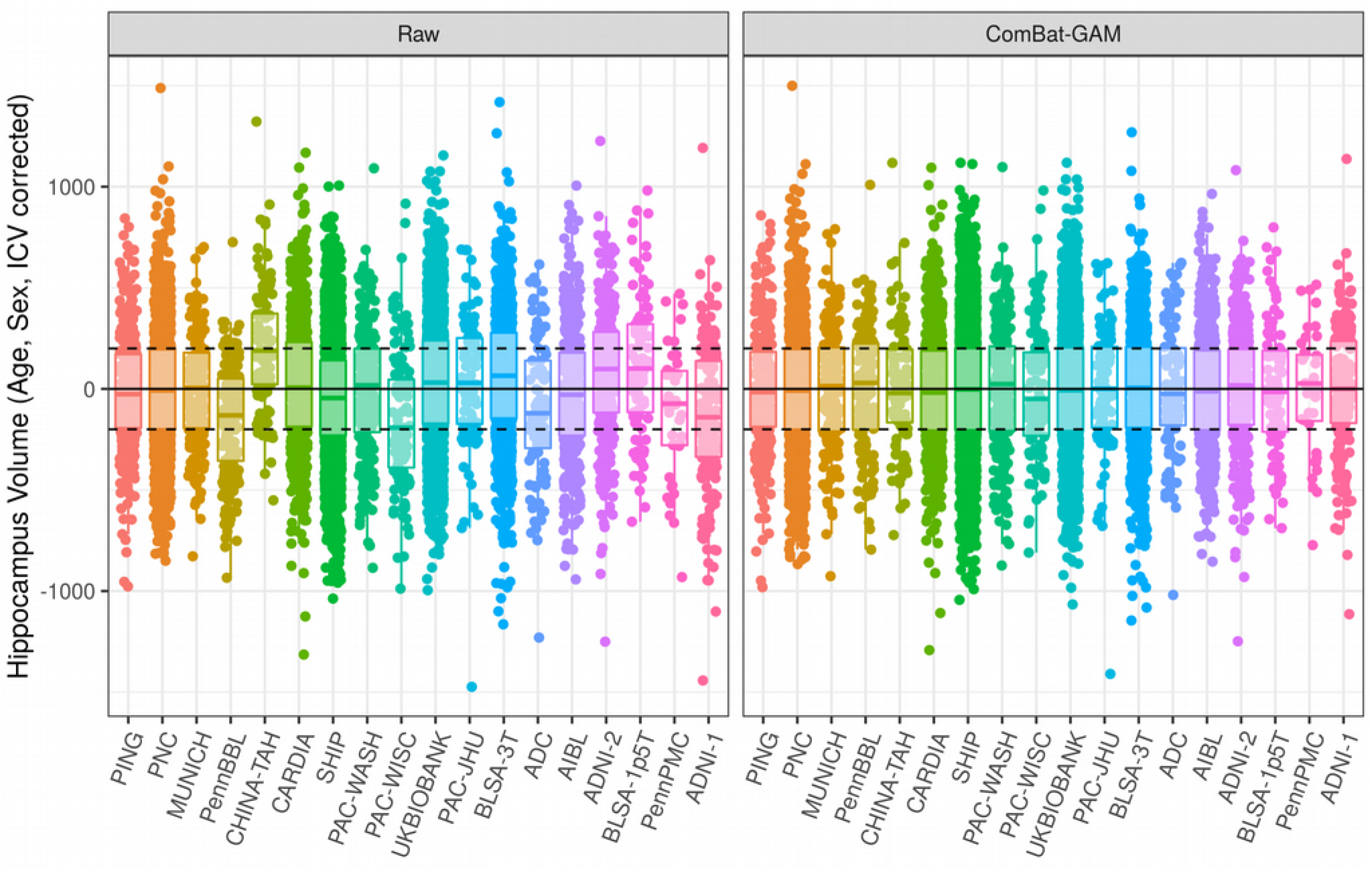
Comparison of hippocampus volumes before and after harmonization, correcting for age, sex, and ICV using GAM. Studies are ordered from youngest to oldest based on median age. In the left panel, volumes were not adjusted for site. In the right panel, volumes were adjusted with ComBat-GAM, which removes location (mean) and scale (variance) differences across sites after controlling for biological covariates. Horizontal lines are plotted at constants at 0, −200, and 200 for visual aid.

Age predictions obtained from the model trained using ROI volumes of participants with 10-fold cross validation were more accurate when the data were harmonized. Figure 7 shows predicted and actual ages for models trained on non-harmonized data, data harmonized with ComBat using a linear age model, and data harmonized using ComBat-GAM. While the application of ComBat with a linear model helped age prediction accuracy compared to no harmonization, the additional use of GAM yielded the best results of the three methods, achieving mean absolute error (MAE) of 5.369.

**Figure 7:**
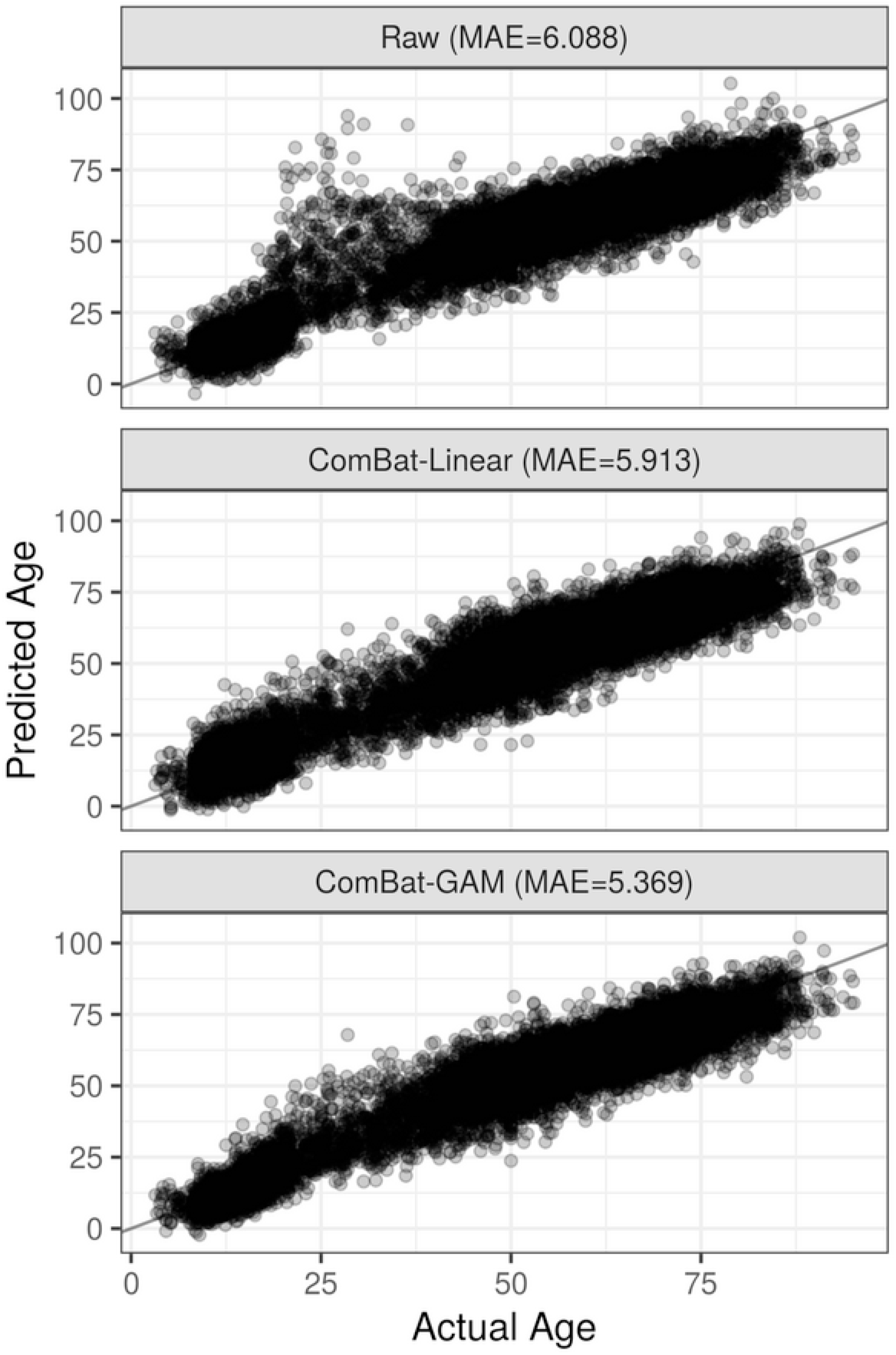
Comparison of age prediction results using three harmonization methods and 10-fold cross validation with a fully-connected neural network using ROI Volumes as input features. MAE is the mean absolute error (i.e. actual age minus predicted age). In the top panel, data were unadjusted for site. In the middle panel, data were harmonized with ComBat using a linear model. In the bottom panel, data were harmonized using ComBat-GAM.

In the leave-site-out validation experiments using the PNC, SHIP, and BLSA-3T as test datasets, harmonization with ComBat-GAM consistently led to improved prediction accuracy for each dataset, compared to using non-harmonized data or using data harmonized with ComBat using a linear age model (Table 3).

**Table 3.** Results of leave-site-out age prediction for each harmonization method.

| Dataset | MAE* obtained for each Harmonization Method |  |  |
| --- | --- | --- | --- |
|  | Raw (non-harmonized) Data | ComBat-Linear | ComBat-GAM |
| PNC (n=1422) | 7.350 | 6.849 | 5.391 |
| SHIP (n=2682) | 6.578 | 6.417 | 6.134 |
| BLSA-3T (n=935) | 6.122 | 6.773 | 5.973 |

*\*MAE: Mean absolute error, i.e. mean absolute difference between predicted and actual ages.*

### 3.4 LIFESPAN age trajectories of ROI volumes

LIFESPAN age trajectories of the third ventricle, hippocampus, thalamus, and occipital pole are presented in Figure 8a and the age trajectories of 4 larger anatomical regions, total gray matter, frontal gray matter, total white matter and deep gray matter, are presented in Figure 8b.

**Figure 8:**
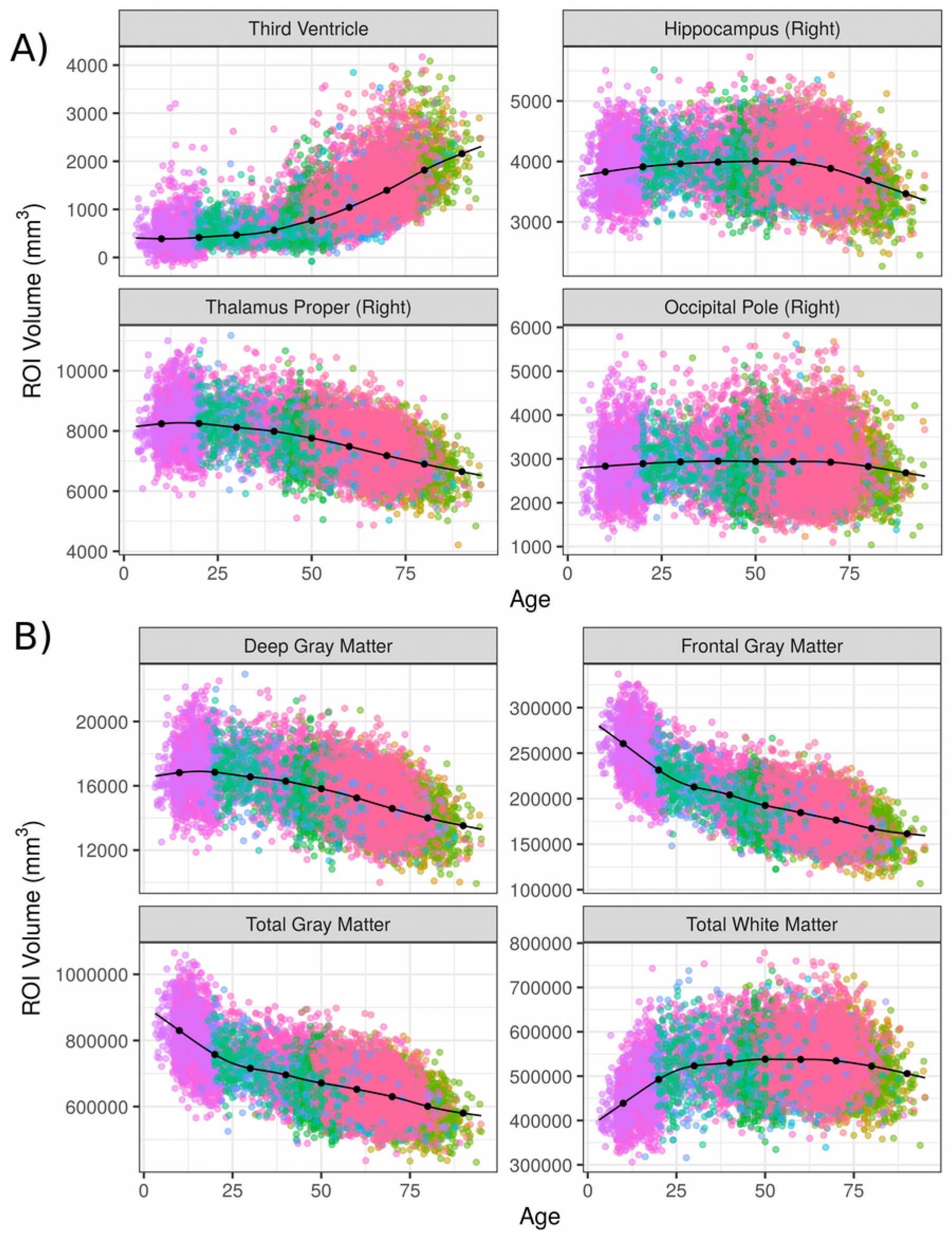
Age trajectories for selected ROI volumes using the combined LIFESPAN dataset with 18 studies spanning the age range 3 – 96. Data were harmonized using ComBat-GAM. The trajectories plotted are for females and assume an average intra-cranial volume (ICV). In A), four ROIs were selected from the set of 145 single ROIs. In B) four ROIs were selected from the set of 113 composite ROIs.

The age trajectories derived from the LIFESPAN data demonstrated variability at both the scales of single ROIs and composite ROIs. At the single ROI level, the hippocampus demonstrated accelerated atrophy late in the lifespan. From age 50 to 60, for example, the percentage difference in hippocampal volume declined by 0.344% over 10 years, according to the age trajectory. In contrast, hippocampal volume declined by 5.132% between age 70 and 80, and by 5.944% from age 80 to 90. Occipital pole volumes were relatively stable throughout the lifespan. Total gray matter volume demonstrated a period of rapid decline during adolescence, followed by gradual decay after age 25. Total white matter volume demonstrated growth during adolescence, stability between ages 30 and 70, and gradual decline after age 75.

Age trajectories for each ROI from the harmonized dataset are made available via a web-based application hosted at the following URL: https://rpomponio.shinyapps.io/neuro_lifespan/. The application allows users to view the age trajectory of any ROI selected from the set of 145 ROIs harmonized in this study, as well as the 113 composite ROIs. The users may upload ROI volumes from a new study to visualize them and compare them with the presented age trajectories. The application also allows users to align their data to pre-calculated trajectories, by removing the location (mean) and scale (variance) differences between new ROI volumes and the reference dataset after controlling for age, sex, and ICV. Figure 9 shows a screenshot of the application being used to visualize the hippocampus volume for an independent dataset together with the LIFESPAN age trajectory for this ROI.

**Figure 9:**
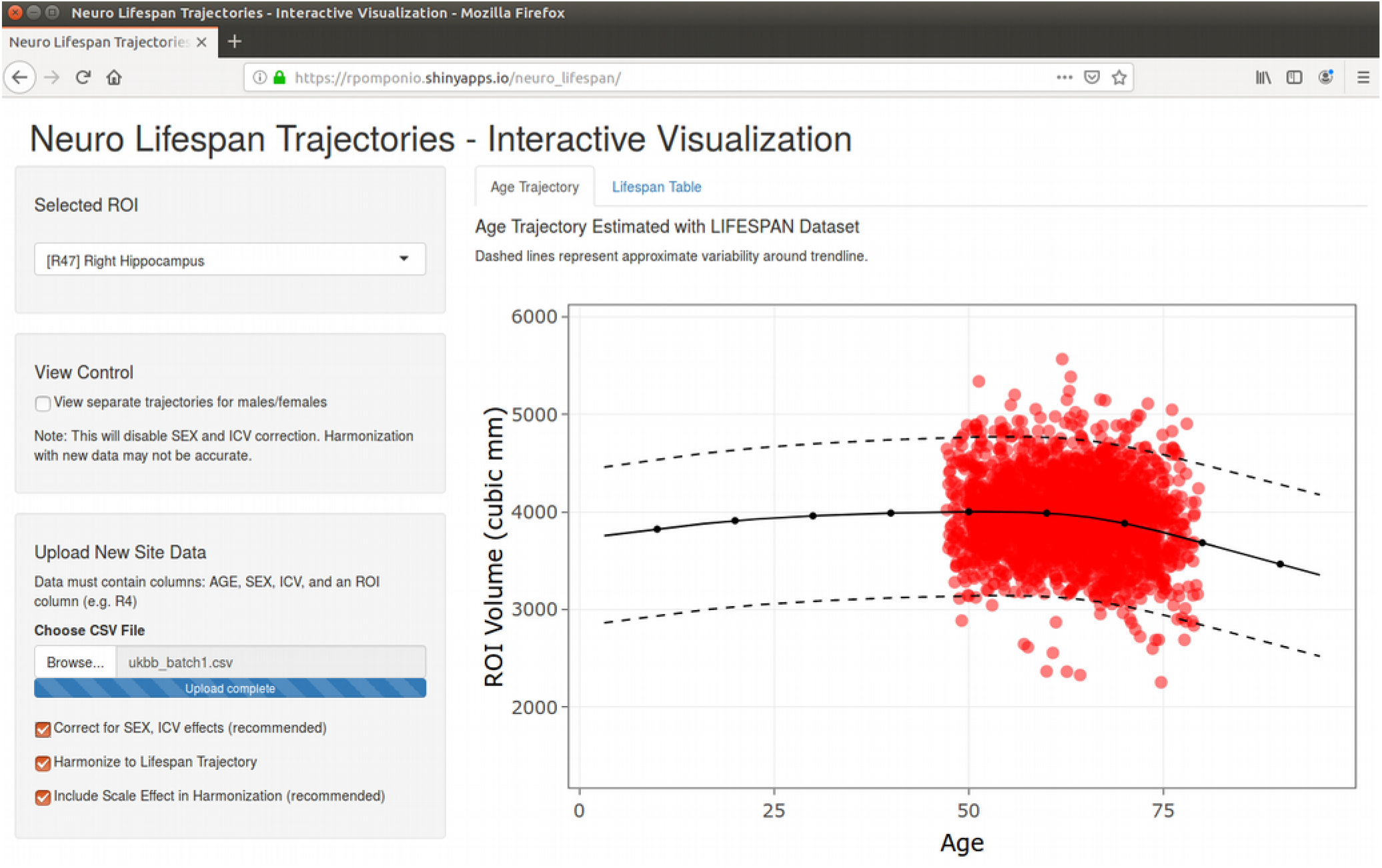
Screenshot of the web-based application that allows visualization of the age trajectory for each anatomical ROI in our dataset. In red, an independent dataset (a supplementary batch of UKBIOBANK) has been uploaded after MUSE segmentation. UKBIOBANK values are aligned to the LIFESPAN age trajectory by removing the location (mean) and scale (variance) differences between new ROI volumes and the reference dataset after controlling for age, sex, and ICV. The application is hosted at the following URL: https://rpomponio.shinyapps.io/neuro_lifespan/.

## 4. Discussion

We described and validated a methodology for harmonization and pooling of neuroimaging data across multiple scanners and cohorts. Using this methodology, as well as regional volumetric measures from 18 neuroimaging studies, we created a large-scale dataset of structural MRI scans covering nearly the entire human lifespan. We applied a fully-automated image processing pipeline to extract regional volumes, followed by an automated quality control procedure to ensure data integrity, and a systematic harmonization method to eliminate site effects while controlling for nonlinear age effects, with the final goal of deriving age trajectories of 258 brain regions at multiple resolution levels. In order to facilitate use of our methodology and data, we developed an interactive visualization and harmonization tool for displaying age trajectories of individual anatomical regions. This tool provides a reference frame for comparing the values of a new cohort against age trajectories estimated from 10,232 participants.

We proposed the use of generalized additive models (GAM) to capture non-linearities in age-related changes in brain structure without over-fitting. Each ROI is modeled by a GAM that includes age as a nonlinear predictor and is optimized via restricted maximum likelihood with regularization to estimate a smooth function. GAMs were previously applied to capture nonlinear trends in a study of brain development in adolescents (Satterthwaite et al., 2014). In our experimental validations using three independent datasets with large sample sizes and spanning different age ranges, we demonstrated that a nonlinear model better captured age-related changes in ROI volumes in different periods of the lifespan compared to linear and quadratic models. The superior performance of GAMs over linear models is consistent with evidence of non-linearity in various anatomical structures, such as gray matter lobes (Giedd et al., 1999), basal ganglia (Ziegler et. al., 2012), and the hippocampus in late-life participants (Allen et al., 2005; Janowitz et al., 2014).

In order to better-understand the behavior of our harmonization procedure relative to the age range covered by each study, we performed simulation experiments leveraging a single-site study in which we introduced artificial site effects. The first conclusion from these simulation experiments was that partially-overlapping age ranges were preferable to disjoint age ranges. This result was expected, as age-disjoint studies should be difficult to harmonize in the presence of nonlinear age effects. The second result suggested that using all available data was preferable to the benefit of balancing across multi-site samples.

Studies have used regional parcellation into anatomical ROIs to understand the brain morphologic changes during the lifespan as well as the effect of disease on the brain (Giedd et al., 1999; Ziegler et al., 2012). Often age has been associated with brain atrophy in various regions (Coffey et al., 1998; Habes et al., 2016), that could be linked to age-related pathologies such as neurodegenerative disorders (Dickerson et al., 2009; Whitwell et al., 2007), but also to the the normal process of aging, which was suggested to be accompanied by demyelination in the white matter and axonal loss (Hinman and Abraham, 2007). The individual’s genetic profile, lifestyle, environment, and disease-related risk factors interact together and contribute to the brain regional vulnerability to age-related changes (Janowitz et al., 2014; Rodrigue et al., 2013). Our harmonized data suggest that there is remarkable variability in the shape and nonlinearity of age trajectories of various ROIs, consistent with previous reports (Courchesne et al., 2000; Walhovd et al. 2011). For example, total gray matter (GM) volume decreases rapidly during late childhood and adolescence, and it continues to decrease, albeit at a much slower rate, in the adult life. We found that total brain white matter (WM) volume follows an inverted-U trajectory, with rapid increases throughout childhood and adolescence then assuming a downward trend around age 60, similar to the trajectory of Cerebral WM volume in Walhovd et al. (2005). Deep GM structures seem to be stable until early adult life, at which point volume declines.

When ROIs are used as building blocks in subsequent analyses, it is important to know the effect of harmonization on subsequently calculated biomarker indices. Toward this goal, we used predicted brain age from a model that summarizes volumetric measures across multiple ROIs as an index that captures the process of typical brain aging, and which has received increasing attention in the literature (Cole and Franke, 2017; Habes et al., 2016; Dosenbach et al., 2010; Erus et al., 2015; Franke et al., 2010). Our results indicated that harmonization has beneficial effects on the calculation of brain age by reducing the prediction error relative to unharmonized data by 11.8%. This is a substantial improvement, especially since it is likely to influence the value of the residuals (brain age – age) that are typically used to flag advanced or resilient brain agers (Eavani et al., 2018).

Our analyses have focused primarily on typically-developing and typically-aging participants, establishing normative age trajectories of brain regions. We included participants without neuropsychiatric disorders; however, to harmonize studies which have a specific neuropsychiatric disease as a focus, data from an appropriate control population is required according to our procedure. Patient data should then follow the same harmonization transformations, but patients should not be used in the calculation of the harmonization model. This is because the underlying assumption behind our approach is that each cohort’s measurements were drawn from the same distribution of values, albeit differing by age, sex, and intra-cranial volume (ICV). Patients with structural brain alterations could violate this assumption and, further, including them in the harmonization would attenuate disease-related effects. Hence, the age trajectory that we provided through the web-interface can serve as a reference based on large control population over a wide age range, and assuming a sufficient control sample is available, could assist with the harmonization task of relatively small pathologic studies, which is otherwise unfeasible.

The current study demonstrates the practical capability of pooling heterogeneous imaging datasets for downstream analysis, particularly at a large scale and in the presence of nonlinear age effects. Future efforts should focus on the application of this framework to new datasets, the inclusion of patient volunteers to derive disease-specific trajectories, and the extension of the current harmonization procedure to longitudinal studies.

## Competing Interests

The authors declare that they have no competing interests.

## Acknowledgements

This work was supported by the National Institute on Aging (grant number 1RF1AG054409) and the National Institute of Mental Health (grant number 5R01MH112070). MH was supported in part by The Allen H. and Selma W. Berkman Charitable Trust (Accelerating Research on Vascular Dementia) and the National Institutes of Health (grant number R01 HL127659-04S1).

