## Supplemental Figures for "Harmonization of large multi-site imaging datasets: Application to 10,232 MRIs for the analysis of imaging patterns of structural brain change throughout the lifespan"

*Supplementary Materials*

**Supplementary Table 1.** Scanner models and acquisition protocol parameters for all studies included in the LIFESPAN dataset.

| ***Dataset*** | ***Scanner*** | ***T1 protocol*** |
|---|---|---|
| ADC | 3T<br>Siemens Trio | MPRAGE<br>1mm x 1mm in-plane<br>1mm slice thickness<br>With 8-channel coil:<br>Flip angle = 15$^{o}$ / TE = 3.87 ms /TR = 1600 ms / TI = 950 ms<br>___________________<br>With 32-channel coil:<br>Flip angle = 9$^{o}$ / TE = 2.89 ms /TR = 1900 ms / TI = 900 ms |
| ADNI-1 | Variable | MPRAGE, saggital<br>FOV: 250 mm<br>256×256 matrix<br>1.2 mm slice thickness<br>Flip angle = 8$^{o}$<br>TE = 3.4 ms<br>TR = 1900 ms |
| ADNI-2 | Variable | MPRAGE<br>208×240×256 matrix<br>1×1×1 mm<br>TE = minimum full echo<br>TR = 2300 ms<br>TI = 900 ms |
| AIBL | 1.5T<br>Siemens Avanto | MPRAGE, sagittal<br>240x256 matrix<br>1mm/1.2 mm slice thickness<br>Flip angle = 9$^{o}$<br>TE = 2.13 ms/2.98&3.05 ms<br>TR = 1900 ms /2300 ms<br>TI = 900 ms |
| BLSA-1.5T | 1.5T<br>GE Signa | SPGR<br>.94×.94×1.5 mm<br>Flip angle = 45$^{o}$<br>TE = 5 ms<br>TR = 35 ms |

| | | |
|---|---|---|
| BLSA-3T | 3T<br>Philips | MPRAGE, sagittal<br>1mm x 1mm in-plane<br>1.2mm slice thickness<br>Flip angle = $8^{o}$<br>TE = 3.1 ms<br>TR = 6.5 ms or 6.8 ms |
| CARDIA | 3T<br>Siemens Tim Trio | MPRAGE, sagittal<br>FOV: 250 mm<br>256x256 matrix<br>1mm slice thickness<br>Flip angle = $9^{o}$<br>TE = 2.89 ms<br>TR = 1900 ms<br>TI = 900 ms |
| CHINA-TAH | 3T<br>GE Discovery MR750 | 3DBRAVO, sagittal<br>Repetition time (ms): 8.156<br>Echo time (ms): 3.172<br>Inversion time (ms): 450<br>Flip angle: 12<br>Number of averages: 1<br>Slice thickness (mm): 1<br>Slice spacing (mm): 1<br>Image columns: 256<br>Image rows: 256<br>Phase encoding direction: ROW<br>Voxel size x (mm): 1<br>Voxel size y (mm): 1<br>Number of volumes: 1<br>Number of slices: 188<br>Number of files: 188<br>Number of frames: 0 |
| MUNICH | | |
| PAC-JHU | 3T<br>Phillips | MPRAGE<br>256×256×170<br>1×1×1.2 mm<br>Flip angle = $8^{o}$<br>TE = 3.1 ms<br>TR = 6.75 ms |
| PAC-WASH | 3T<br>Siemens Tim Trio | 256×256×176 matrix<br>1×1×1 mm<br>Flip angle = $8^{o}$<br>TE = 3.16 ms<br>TR = 2400 ms<br>TI = 1000 ms |

| | | |
|---|---|---|
| PAC-WISC | 3T<br>GE Discovery MR750 (Waukesha, WI, USA) with 8-channel head coil and parallel imaging with Array Spatial Sensitivity Encoding Technique | SPGR, axial<br>FOV: 256 mm<br>56x256 matrix<br>1mm slice thickness<br>Flip angle = $12^{o}$<br>TR = 8.1 ms<br>TI = 450 ms<br>TE = 3.2 ms |
| PennBBL | | |
| PennPMC | 3T<br>Siemens Trio | MPRAGE<br>1mm x 1mm in-plane<br>1mm slice thickness<br>With 8-channel coil:<br>Flip angle = $15^{o}$ / TE = 3.87 ms /TR = 1600 ms / TI = 950 ms<br>____________________<br>With 32-channel coil:<br>Flip angle = $9^{o}$ / TE = 2.89 ms /TR = 1900 ms / TI = 900 ms |
| PING | 3T<br>Siemens TrioTim<br>Philips Achieva<br>GE Signa<br>GE Discovery | MPRAGE<br>TR = Variable, 6.8ms – 8.1ms<br>TE = Variable, 3.1ms – 4.33ms<br>TI = Variable, 640ms – 1100ms<br>Flip Angle = Variable, $7^{o}$ - $8^{o}$<br>Matrix size = Variable<br>Voxel Size = 1 x 1 x 1.2<br>Acquisition Time = Variable, 8:05 – 9:19.7 |
| PNC | 3T<br>Siemens Tim Trio | MPRAGE, axial<br>1mm slice thickness<br>Flip angle =$9^{o}$<br>TR = 1810 ms<br>TI = 1100 ms |
| SHIP | 1.5T<br>Siemens Magnetom Avanto | MPRAGE, axial<br>1mm x 1mm in-plane<br>1mm slice thickness<br>Flip angle = $15^{o}$<br>TE = 34 ms<br>TR = 19000 ms |
| UKBIOBANK | 3T<br>Siemens Skyra (VD13) | MPRAGE, sagittal<br>1mm x 1mm in-plane<br>1mm slice thickness<br>TR = 2000 ms<br>TI = 880 ms |

**Supplementary Table 2.** Regions of Interest (ROI) used to obtain volumetric measures in the LIFESPAN dataset.

| ROI_INDEX | ROI_NAME | ROI_RESOLUTION |
|---|---|---|
| 4 | 3rd Ventricle | Single |
| 11 | 4th Ventricle | Single |
| 23 | Right Accumbens Area | Single |
| 30 | Left Accumbens Area | Single |
| 31 | Right Amygdala | Single |
| 32 | Left Amygdala | Single |
| 35 | Brain Stem | Single |
| 36 | Right Caudate | Single |
| 37 | Left Caudate | Single |
| 38 | Right Cerebellum Exterior | Single |
| 39 | Left Cerebellum Exterior | Single |
| 40 | Right Cerebellum White Matter | Single |
| 41 | Left Cerebellum White Matter | Single |
| 47 | Right Hippocampus | Single |
| 48 | Left Hippocampus | Single |
| 49 | Right Inf Lat Vent | Single |
| 50 | Left Inf Lat Vent | Single |
| 51 | Right Lateral Ventricle | Single |
| 52 | Left Lateral Ventricle | Single |
| 55 | Right Pallidum | Single |
| 56 | Left Pallidum | Single |
| 57 | Right Putamen | Single |
| 58 | Left Putamen | Single |
| 59 | Right Thalamus Proper | Single |
| 60 | Left Thalamus Proper | Single |
| 61 | Right Ventral DC | Single |
| 62 | Left Ventral DC | Single |
| 71 | Cerebellar Vermal Lobules I-V | Single |
| 72 | Cerebellar Vermal Lobules VI-VII | Single |
| 73 | Cerebellar Vermal Lobules VIII-X | Single |
| 75 | Left Basal Forebrain | Single |
| 76 | Right Basal Forebrain | Single |
| 81 | frontal lobe WM right | Single |
| 82 | frontal lobe WM left | Single |
| 83 | occipital lobe WM right | Single |
| 84 | occipital lobe WM left | Single |
| 85 | parietal lobe WM right | Single |
| 86 | parietal lobe WM left | Single |
| 87 | temporal lobe WM right | Single |
| 88 | temporal lobe WM left | Single |
| 89 | fornix right | Single |
| 90 | fornix left | Single |
| 91 | anterior limb of internal capsule right | Single |
| 92 | anterior limb of internal capsule left | Single |
| 93 | posterior limb of internal capsule inc. cerebral peduncle right | Single |
| 94 | posterior limb of internal capsule inc. cerebral peduncle left | Single |
| 95 | corpus callosum | Single |

| | | |
|---|---|---|
| 100 | Right ACgG anterior cingulate gyrus | Single |
| 101 | Left ACgG anterior cingulate gyrus | Single |
| 102 | Right AIns anterior insula | Single |
| 103 | Left AIns anterior insula | Single |
| 104 | Right AOrG anterior orbital gyrus | Single |
| 105 | Left AOrG anterior orbital gyrus | Single |
| 106 | Right AnG angular gyrus | Single |
| 107 | Left AnG angular gyrus | Single |
| 108 | Right Calc calcarine cortex | Single |
| 109 | Left Calc calcarine cortex | Single |
| 112 | Right CO central operculum | Single |
| 113 | Left CO central operculum | Single |
| 114 | Right Cun cuneus | Single |
| 115 | Left Cun cuneus | Single |
| 116 | Right Ent entorhinal area | Single |
| 117 | Left Ent entorhinal area | Single |
| 118 | Right FO frontal operculum | Single |
| 119 | Left FO frontal operculum | Single |
| 120 | Right FRP frontal pole | Single |
| 121 | Left FRP frontal pole | Single |
| 122 | Right FuG fusiform gyrus | Single |
| 123 | Left FuG fusiform gyrus | Single |
| 124 | Right GRe gyrus rectus | Single |
| 125 | Left GRe gyrus rectus | Single |
| 128 | Right IOG inferior occipital gyrus | Single |
| 129 | Left IOG inferior occipital gyrus | Single |
| 132 | Right ITG inferior temporal gyrus | Single |
| 133 | Left ITG inferior temporal gyrus | Single |
| 134 | Right LiG lingual gyrus | Single |
| 135 | Left LiG lingual gyrus | Single |
| 136 | Right LOrG lateral orbital gyrus | Single |
| 137 | Left LOrG lateral orbital gyrus | Single |
| 138 | Right MCgG middle cingulate gyrus | Single |
| 139 | Left MCgG middle cingulate gyrus | Single |
| 140 | Right MFC medial frontal cortex | Single |
| 141 | Left MFC medial frontal cortex | Single |
| 142 | Right MFG middle frontal gyrus | Single |
| 143 | Left MFG middle frontal gyrus | Single |
| 144 | Right MOG middle occipital gyrus | Single |
| 145 | Left MOG middle occipital gyrus | Single |
| 146 | Right MOrG medial orbital gyrus | Single |
| 147 | Left MOrG medial orbital gyrus | Single |
| 148 | Right MPoG postcentral gyrus medial segment | Single |
| 149 | Left MPoG postcentral gyrus medial segment | Single |
| 150 | Right MPrG precentral gyrus medial segment | Single |
| 151 | Left MPrG precentral gyrus medial segment | Single |
| 152 | Right MSFG superior frontal gyrus medial segment | Single |
| 153 | Left MSFG superior frontal gyrus medial segment | Single |
| 154 | Right MTG middle temporal gyrus | Single |
| 155 | Left MTG middle temporal gyrus | Single |
| 156 | Right OCP occipital pole | Single |
| 157 | Left OCP occipital pole | Single |
| 160 | Right OFuG occipital fusiform gyrus | Single |

| | |
|---|---|
| 161 Left OFuG occipital fusiform gyrus | Single |
| 162 Right OpIFG opercular part of the inferior frontal gyrus | Single |
| 163 Left OpIFG opercular part of the inferior frontal gyrus | Single |
| 164 Right OrIFG orbital part of the inferior frontal gyrus | Single |
| 165 Left OrIFG orbital part of the inferior frontal gyrus | Single |
| 166 Right PCgG posterior cingulate gyrus | Single |
| 167 Left PCgG posterior cingulate gyrus | Single |
| 168 Right PCu precuneus | Single |
| 169 Left PCu precuneus | Single |
| 170 Right PHG parahippocampal gyrus | Single |
| 171 Left PHG parahippocampal gyrus | Single |
| 172 Right PIns posterior insula | Single |
| 173 Left PIns posterior insula | Single |
| 174 Right PO parietal operculum | Single |
| 175 Left PO parietal operculum | Single |
| 176 Right PoG postcentral gyrus | Single |
| 177 Left PoG postcentral gyrus | Single |
| 178 Right POrG posterior orbital gyrus | Single |
| 179 Left POrG posterior orbital gyrus | Single |
| 180 Right PP planum polare | Single |
| 181 Left PP planum polare | Single |
| 182 Right PrG precentral gyrus | Single |
| 183 Left PrG precentral gyrus | Single |
| 184 Right PT planum temporale | Single |
| 185 Left PT planum temporale | Single |
| 186 Right SCA subcallosal area | Single |
| 187 Left SCA subcallosal area | Single |
| 190 Right SFG superior frontal gyrus | Single |
| 191 Left SFG superior frontal gyrus | Single |
| 192 Right SMC supplementary motor cortex | Single |
| 193 Left SMC supplementary motor cortex | Single |
| 194 Right SMG supramarginal gyrus | Single |
| 195 Left SMG supramarginal gyrus | Single |
| 196 Right SOG superior occipital gyrus | Single |
| 197 Left SOG superior occipital gyrus | Single |
| 198 Right SPL superior parietal lobule | Single |
| 199 Left SPL superior parietal lobule | Single |
| 200 Right STG superior temporal gyrus | Single |
| 201 Left STG superior temporal gyrus | Single |
| 202 Right TMP temporal pole | Single |
| 203 Left TMP temporal pole | Single |
| 204 Right TrIFG triangular part of the inferior frontal gyrus | Single |
| 205 Left TrIFG triangular part of the inferior frontal gyrus | Single |
| 206 Right TTG transverse temporal gyrus | Single |
| 207 Left TTG transverse temporal gyrus | Single |
| 301 FRONTAL_INFERIOR_GM | Composite |
| 302 FRONTAL_INSULAR_GM | Composite |
| 303 FRONTAL_LATERAL_GM | Composite |
| 304 FRONTAL_MEDIAL_GM | Composite |
| 305 FRONTAL_OPERCULAR_GM | Composite |
| 306 LIMBIC_CINGULATE_GM | Composite |
| 307 LIMBIC_MEDIALTEMPORAL_GM | Composite |
| 308 OCCIPITAL_INFERIOR_GM | Composite |

| | | |
|---|---|---|
| 309 | OCCIPITAL_LATERAL_GM | Composite |
| 310 | OCCIPITAL_MEDIAL_GM | Composite |
| 311 | PARIETAL_LATERAL_GM | Composite |
| 312 | PARIETAL_MEDIAL_GM | Composite |
| 313 | TEMPORAL_INFERIOR_GM | Composite |
| 314 | TEMPORAL_LATERAL_GM | Composite |
| 315 | TEMPORAL_SUPRATEMPORAL_GM | Composite |
| 316 | FRONTAL_INFERIOR_GM_L | Composite |
| 317 | FRONTAL_INSULAR_GM_L | Composite |
| 318 | FRONTAL_LATERAL_GM_L | Composite |
| 319 | FRONTAL_MEDIAL_GM_L | Composite |
| 320 | FRONTAL_OPERCULAR_GM_L | Composite |
| 321 | LIMBIC_CINGULATE_GM_L | Composite |
| 322 | LIMBIC_MEDIALTEMPORAL_GM_L | Composite |
| 323 | OCCIPITAL_INFERIOR_GM_L | Composite |
| 324 | OCCIPITAL_LATERAL_GM_L | Composite |
| 325 | OCCIPITAL_MEDIAL_GM_L | Composite |
| 326 | PARIETAL_LATERAL_GM_L | Composite |
| 327 | PARIETAL_MEDIAL_GM_L | Composite |
| 328 | TEMPORAL_INFERIOR_GM_L | Composite |
| 329 | TEMPORAL_LATERAL_GM_L | Composite |
| 330 | TEMPORAL_SUPRATEMPORAL_GM_L | Composite |
| 331 | FRONTAL_INFERIOR_GM_R | Composite |
| 332 | FRONTAL_INSULAR_GM_R | Composite |
| 333 | FRONTAL_LATERAL_GM_R | Composite |
| 334 | FRONTAL_MEDIAL_GM_R | Composite |
| 335 | FRONTAL_OPERCULAR_GM_R | Composite |
| 336 | LIMBIC_CINGULATE_GM_R | Composite |
| 337 | LIMBIC_MEDIALTEMPORAL_GM_R | Composite |
| 338 | OCCIPITAL_INFERIOR_GM_R | Composite |
| 339 | OCCIPITAL_LATERAL_GM_R | Composite |
| 340 | OCCIPITAL_MEDIAL_GM_R | Composite |
| 341 | PARIETAL_LATERAL_GM_R | Composite |
| 342 | PARIETAL_MEDIAL_GM_R | Composite |
| 343 | TEMPORAL_INFERIOR_GM_R | Composite |
| 344 | TEMPORAL_LATERAL_GM_R | Composite |
| 345 | TEMPORAL_SUPRATEMPORAL_GM_R | Composite |
| 401 | BASAL_GANGLIA | Composite |
| 402 | DEEP_GM | Composite |
| 403 | DEEP_WM | Composite |
| 404 | FRONTAL_GM | Composite |
| 405 | FRONTAL_WM | Composite |
| 406 | LIMBIC_GM | Composite |
| 407 | OCCIPITAL_GM | Composite |
| 408 | OCCIPITAL_WM | Composite |
| 409 | PARIETAL_GM | Composite |
| 410 | PARIETAL_WM | Composite |
| 411 | TEMPORAL_GM | Composite |
| 412 | TEMPORAL_WM | Composite |
| 413 | BASAL_GANGLIA_L | Composite |
| 414 | DEEP_GM_L | Composite |
| 415 | DEEP_WM_L | Composite |
| 416 | FRONTAL_GM_L | Composite |

| | | |
|---|---|---|
| 417 | FRONTAL_WM_L | Composite |
| 418 | LIMBIC_GM_L | Composite |
| 419 | OCCIPITAL_GM_L | Composite |
| 420 | OCCIPITAL_WM_L | Composite |
| 421 | PARIETAL_GM_L | Composite |
| 422 | PARIETAL_WM_L | Composite |
| 423 | TEMPORAL_GM_L | Composite |
| 424 | TEMPORAL_WM_L | Composite |
| 425 | BASAL_GANGLIA_R | Composite |
| 426 | DEEP_GM_R | Composite |
| 427 | DEEP_WM_R | Composite |
| 428 | FRONTAL_GM_R | Composite |
| 429 | FRONTAL_WM_R | Composite |
| 430 | LIMBIC_GM_R | Composite |
| 431 | OCCIPITAL_GM_R | Composite |
| 432 | OCCIPITAL_WM_R | Composite |
| 433 | PARIETAL_GM_R | Composite |
| 434 | PARIETAL_WM_R | Composite |
| 435 | TEMPORAL_GM_R | Composite |
| 436 | TEMPORAL_WM_R | Composite |
| 501 | CORPUS_CALLOSUM | Composite |
| 502 | CEREBELLUM | Composite |
| 503 | DEEP_WM_GM | Composite |
| 504 | FRONTAL | Composite |
| 505 | LIMBIC | Composite |
| 506 | OCCIPITAL | Composite |
| 507 | PARIETAL | Composite |
| 508 | TEMPORAL | Composite |
| 509 | VENTRICLE | Composite |
| 510 | CEREBELLUM_L | Composite |
| 511 | DEEP_WM_GM_L | Composite |
| 512 | FRONTAL_L | Composite |
| 513 | LIMBIC_L | Composite |
| 514 | OCCIPITAL_L | Composite |
| 515 | PARIETAL_L | Composite |
| 516 | TEMPORAL_L | Composite |
| 517 | VENTRICLE_L | Composite |
| 518 | CEREBELLUM_R | Composite |
| 519 | DEEP_WM_GM_R | Composite |
| 520 | FRONTAL_R | Composite |
| 521 | LIMBIC_R | Composite |
| 522 | OCCIPITAL_R | Composite |
| 523 | PARIETAL_R | Composite |
| 524 | TEMPORAL_R | Composite |
| 525 | VENTRICLE_R | Composite |
| 601 | GM | Composite |
| 604 | WM | Composite |
| 606 | GM_L | Composite |
| 607 | WM_L | Composite |
| 613 | GM_R | Composite |
| 614 | WM_R | Composite |
| 701 | TOTALBRAIN | Composite |
| 702 | ICV | Composite |

**Supplementary Table 3.** Anatomical regions used to identify outliers in the quality control (QC) procedure.

| ROI_INDEX | ROI_NAME |
|---|---|
| 23 | Right Accumbens Area |
| 30 | Left Accumbens Area |
| 31 | Right Amygdala |
| 32 | Left Amygdala |
| 36 | Right Caudate |
| 37 | Left Caudate |
| 38 | Right Cerebellum Exterior |
| 39 | Left Cerebellum Exterior |
| 40 | Right Cerebellum White Matter |
| 41 | Left Cerebellum White Matter |
| 47 | Right Hippocampus |
| 48 | Left Hippocampus |
| 55 | Right Pallidum |
| 56 | Left Pallidum |
| 57 | Right Putamen |
| 58 | Left Putamen |
| 59 | Right Thalamus Proper |
| 60 | Left Thalamus Proper |
| 71 | Cerebellar Vermal Lobules I-V |
| 72 | Cerebellar Vermal Lobules VI-VII |
| 73 | Cerebellar Vermal Lobules VIII-X |
| 75 | Left Basal Forebrain |
| 76 | Right Basal Forebrain |
| 81 | frontal lobe WM right |
| 82 | frontal lobe WM left |
| 83 | occipital lobe WM right |
| 84 | occipital lobe WM left |
| 85 | parietal lobe WM right |
| 86 | parietal lobe WM left |
| 87 | temporal lobe WM right |
| 88 | temporal lobe WM left |
| 89 | fornix right |
| 90 | fornix left |
| 91 | anterior limb of internal capsule right |
| 92 | anterior limb of internal capsule left |
| 93 | posterior limb of internal capsule inc. cerebral peduncle right |
| 94 | posterior limb of internal capsule inc. cerebral peduncle left |
| 95 | corpus callosum |
| 316 | FRONTAL_INFERIOR_GM_L |
| 317 | FRONTAL_INSULAR_GM_L |
| 318 | FRONTAL_LATERAL_GM_L |
| 319 | FRONTAL_MEDIAL_GM_L |
| 320 | FRONTAL_OPERCULAR_GM_L |
| 321 | LIMBIC_CINGULATE_GM_L |
| 322 | LIMBIC_MEDIALTEMPORAL_GM_L |
| 323 | OCCIPITAL_INFERIOR_GM_L |
| 324 | OCCIPITAL_LATERAL_GM_L |
| 325 | OCCIPITAL_MEDIAL_GM_L |

326 PARIETAL_LATERAL_GM_L
327 PARIETAL_MEDIAL_GM_L
328 TEMPORAL_INFERIOR_GM_L
329 TEMPORAL_LATERAL_GM_L
330 TEMPORAL_SUPRATEMPORAL_GM_L
331 FRONTAL_INFERIOR_GM_R
332 FRONTAL_INSULAR_GM_R
333 FRONTAL_LATERAL_GM_R
334 FRONTAL_MEDIAL_GM_R
335 FRONTAL_OPERCULAR_GM_R
336 LIMBIC_CINGULATE_GM_R
337 LIMBIC_MEDIALTEMPORAL_GM_R
338 OCCIPITAL_INFERIOR_GM_R
339 OCCIPITAL_LATERAL_GM_R
340 OCCIPITAL_MEDIAL_GM_R
341 PARIETAL_LATERAL_GM_R
342 PARIETAL_MEDIAL_GM_R
343 TEMPORAL_INFERIOR_GM_R
344 TEMPORAL_LATERAL_GM_R
345 TEMPORAL_SUPRATEMPORAL_GM_R
702 ICV

**Supplementary Table 4.** Summary of scans identified and excluded as outliers in LIFESPAN dataset.

| *Dataset* | *Original No. Subjects* | *No. Outlier Subjects (%)* | *No. Female Outliers (%)* | *Outliers Age Range* |
|---|---|---|---|---|
| ADC | 104 | 1 (1.0) | 0 (0.0) | [83, 83] |
| ADNI-1 | 189 | 7 (3.7) | 5 (71.4) | [72, 88] |
| ADNI-2 | 324 | 12 (3.7) | 9 (75.0) | [63, 86] |
| AIBL | 446 | 17 (3.8) | 7 (41.2) | [63, 88] |
| BLSA-1.5T | 92 | 1 (1.1) | 0 (0.0) | [69, 69] |
| BLSA-3T | 964 | 29 (3.0) | 20 (69.0) | [22, 91] |
| CARDIA | 719 | 26 (3.6) | 17 (65.4) | [43, 56] |
| CHINA-TAH | 105 | 0 (0.0) | 0 (0.0) | N/A |
| MUNICH | 173 | 4 (2.3) | 2 (50.0) | [30, 54] |
| PAC-JHU | 94 | 2 (2.1) | 2 (1.0) | [72, 74] |
| PAC-WASH | 247 | 8 (3.2) | 4 (50.0) | [47, 77] |
| PAC-WISC | 127 | 5 (3.9) | 3 (60.0) | [52, 71] |
| PennBBL | 170 | 6 (3.5) | 5 (83.3) | [22, 86] |
| PennPMC | 41 | 1 (2.4) | 1 (1.0) | [65, 65] |
| PING | 306 | 5 (1.6) | 3 (60.0) | [4, 21] |
| PNC | 1445 | 23 (1.6) | 13 (56.5) | [8, 22] |
| SHIP | 2739 | 57 (2.1) | 36 (63.2) | [21, 90] |
| UKBIOBANK | 2201 | 50 (2.3) | 30 (60.0) | [49, 80] |

**Supplementary Figure 1.** Comparison of selected ROI volumes before and after harmonization, correcting for age, sex, and ICV using a generalized additive model (GAM).

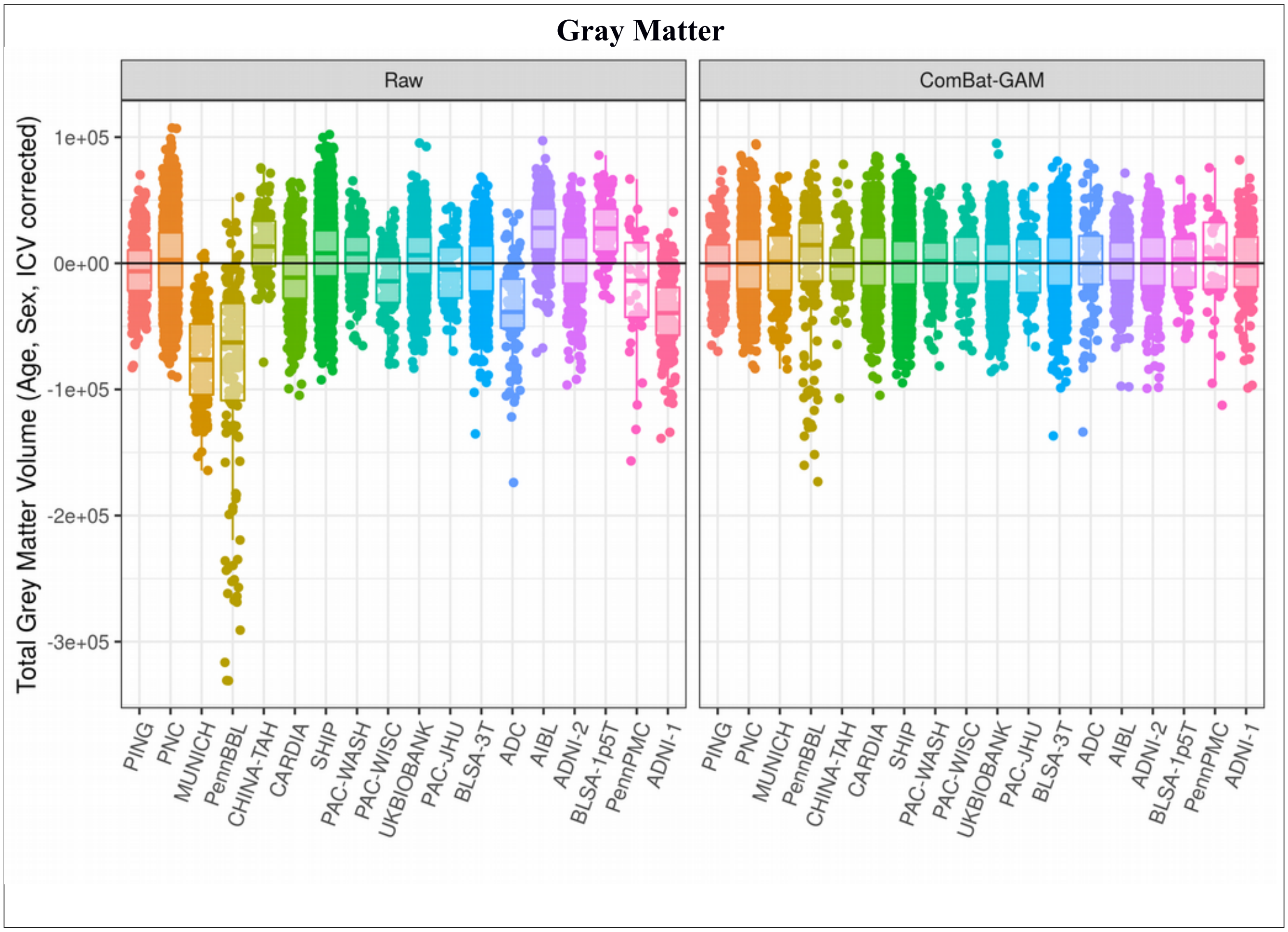

**White Matter**

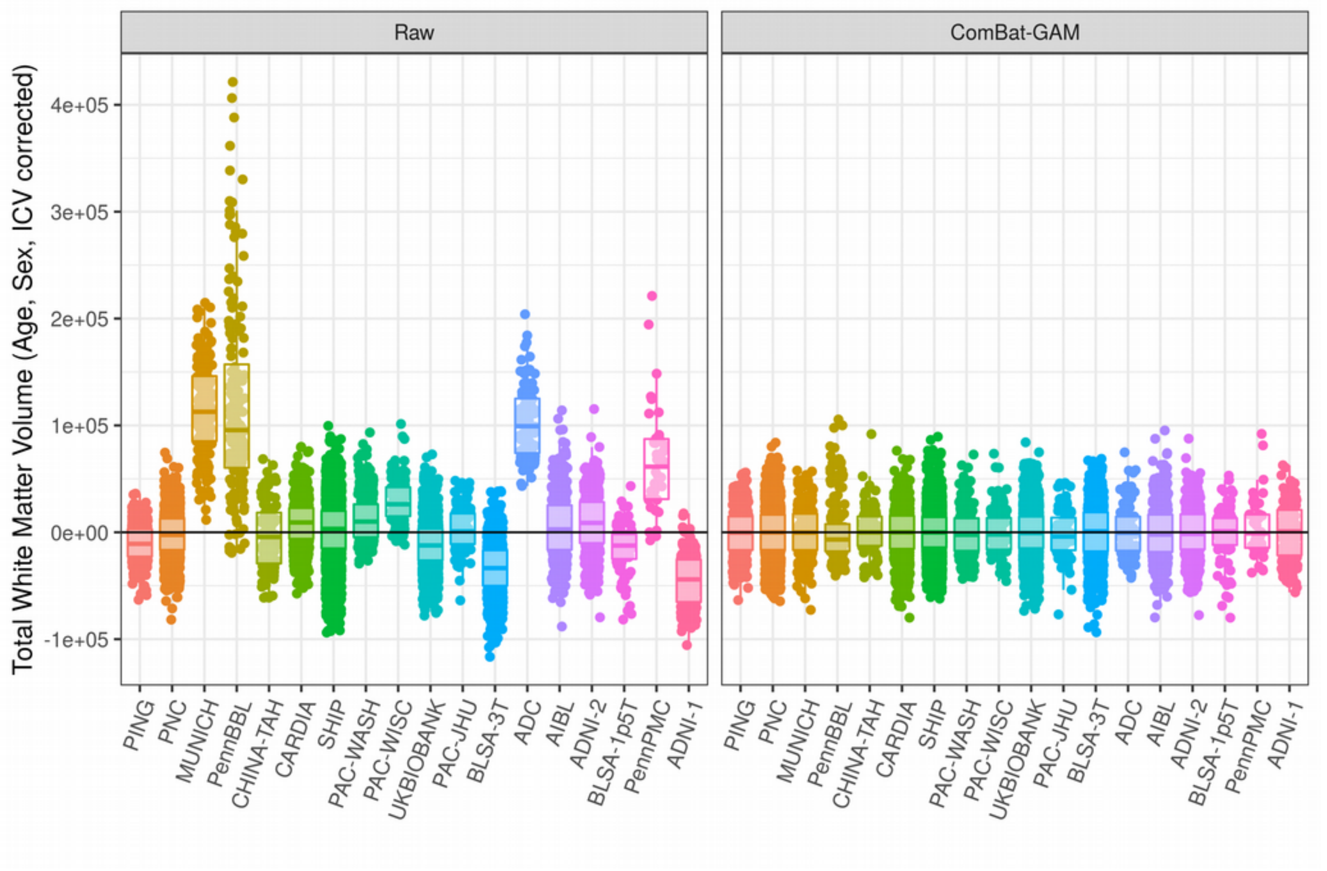

**Amygdala**

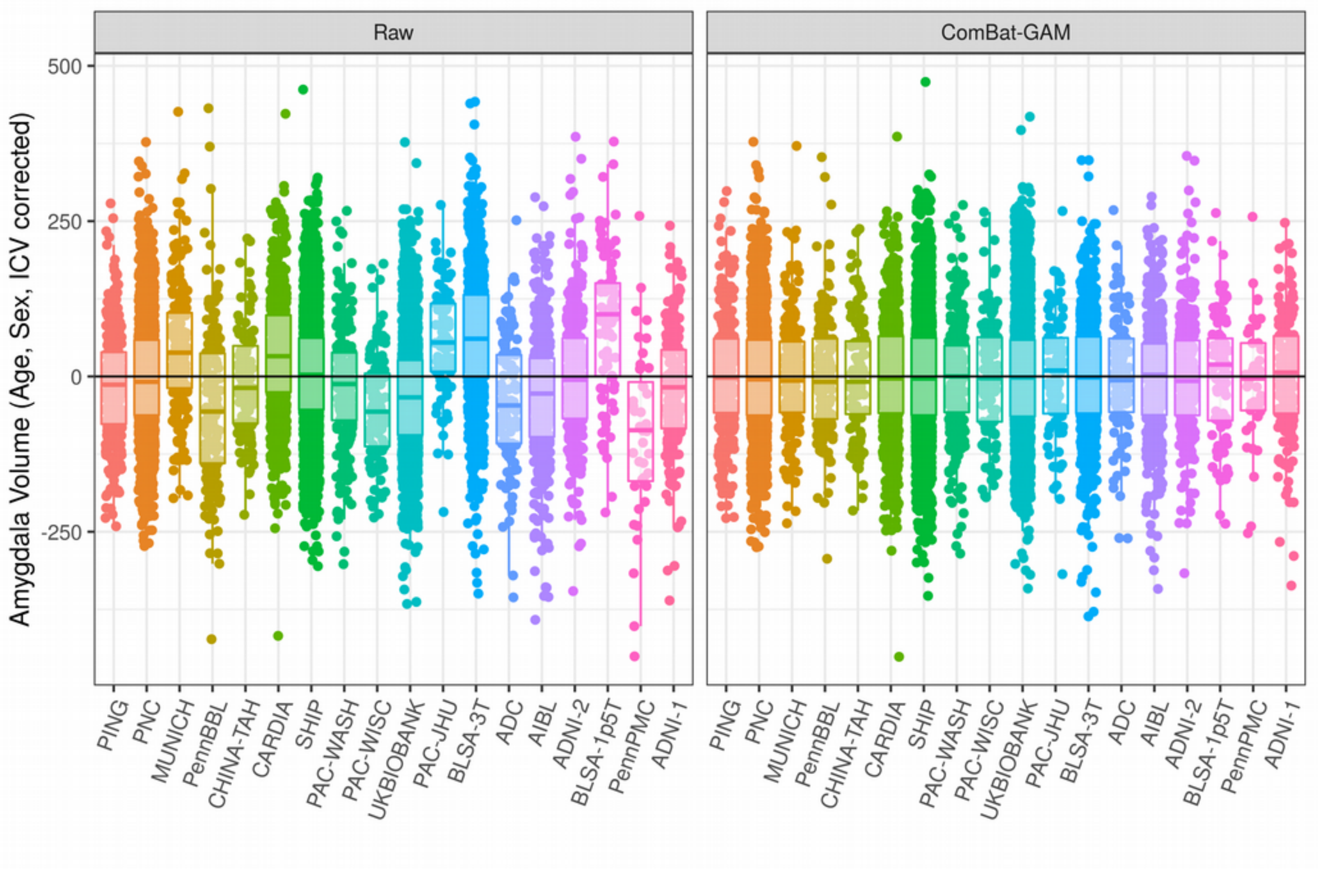
Raw
ComBat-GAM
Amygdala Volume (Age, Sex, ICV corrected)
500
250
0
-250
PING
PNC
MUNICH
PennBBL
CHINA-TAH
CARDIA
SHIP
PAC-WASH
PAC-WISC
UKBIOBANK
PAC-JHU
BLSA-3T
ADC
AIBL
ADNI-2
BLSA-1p5T
PennPMC
ADNI-1

### Thalamus

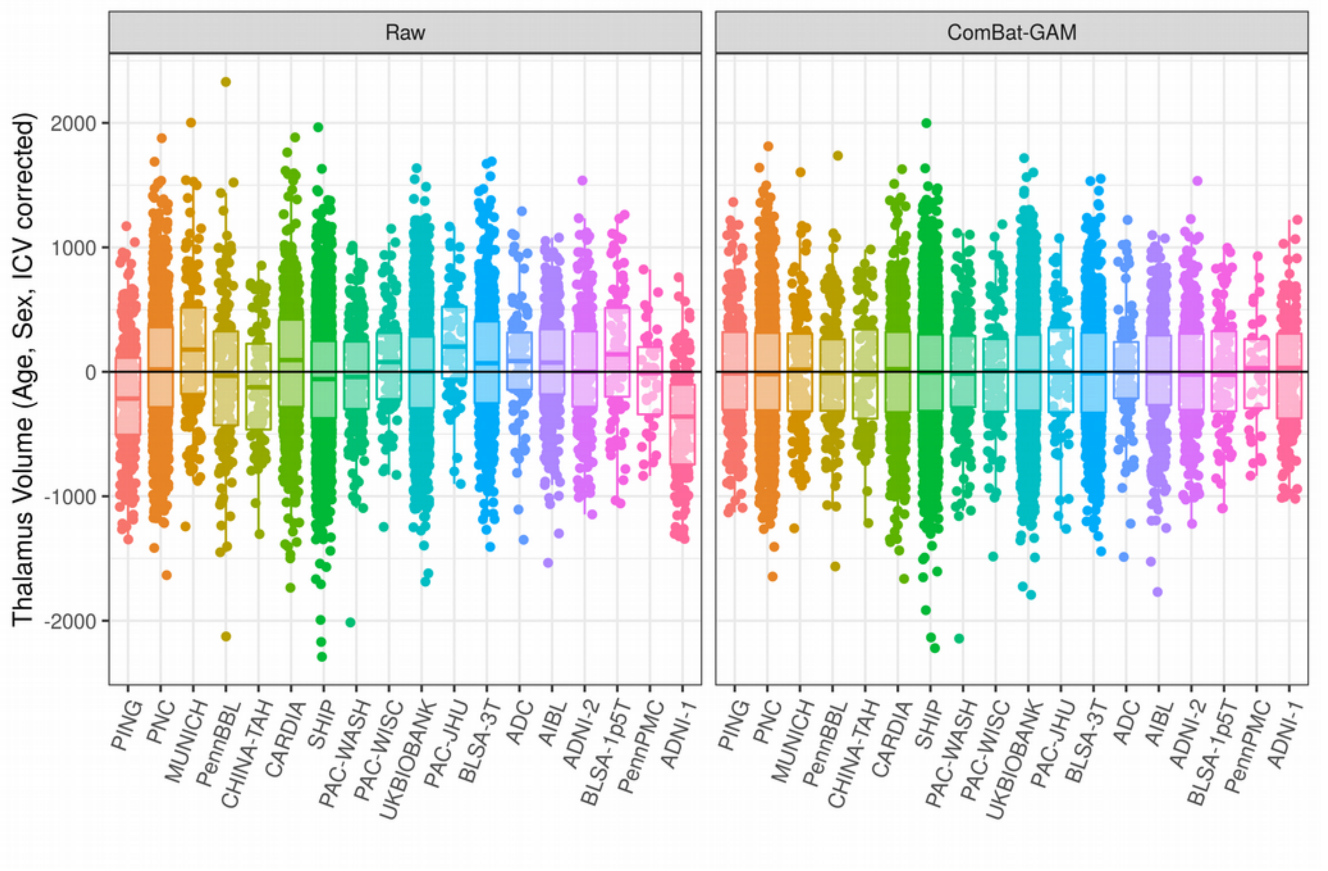

Raw
ComBat-GAM
Thalamus Volume (Age, Sex, ICV corrected)
2000
1000
0
-1000
-2000
PING
PNC
MUNICH
PennBBL
CHINA-TAH
CARDIA
SHIP
PAC-WASH
PAC-WISC
UKBIOBANK
PAC-JHU
BLSA-3T
ADC
AIBL
ADNI-2
BLSA-1p5T
PennPMC
ADNI-1
